# Nanomolar inhibition of SARS-CoV-2 infection by an unmodified peptide targeting the pre-hairpin intermediate of the spike protein

**DOI:** 10.1101/2022.08.11.503553

**Authors:** Kailu Yang, Chuchu Wang, Alex J. B. Kreutzberger, Ravi Ojha, Suvi Kuivanen, Sergio Couoh-Cardel, Serena Muratcioglu, Timothy J. Eisen, K. Ian White, Richard G. Held, Subu Subramanian, Kendra Marcus, Richard A. Pfuetzner, Luis Esquivies, Catherine A. Doyle, John Kuriyan, Olli Vapalahti, Giuseppe Balistreri, Tomas Kirchhausen, Axel T. Brunger

## Abstract

Variants of severe acute respiratory syndrome coronavirus 2 (SARS-CoV-2) challenge currently available COVID-19 vaccines and monoclonal antibody therapies through epitope change on the receptor binding domain of the viral spike glycoprotein. Hence, there is a specific urgent need for alternative antivirals that target processes less likely to be affected by mutation, such as the membrane fusion step of viral entry into the host cell. One such antiviral class includes peptide inhibitors which block formation of the so-called HR1HR2 six-helix bundle of the SARS-CoV-2 spike (S) protein and thus interfere with viral membrane fusion. Here we performed structural studies of the HR1HR2 bundle, revealing an extended, well-folded N-terminal region of HR2 that interacts with the HR1 triple helix. Based on this structure, we designed an extended HR2 peptide that achieves single-digit nanomolar inhibition of SARS-CoV-2 in cell-based fusion, VSV-SARS-CoV-2 chimera, and authentic SARS-CoV-2 infection assays without the need for modifications such as lipidation or chemical stapling. The peptide also strongly inhibits all major SARS-CoV-2 variants to date. This extended peptide is ~100-fold more potent than all previously published short, unmodified HR2 peptides, and it has a very long inhibition lifetime after washout in virus infection assays, suggesting that it targets a pre-hairpin intermediate of the SARS-CoV-2 S protein. Together, these results suggest that regions outside the HR2 helical region may offer new opportunities for potent peptide-derived therapeutics for SARS-CoV-2 and its variants, and even more distantly related viruses, and provide further support for the pre-hairpin intermediate of the S protein.

**Significance Statement:** SARS-CoV-2 infection requires fusion of viral and host membranes, mediated by the viral spike glycoprotein (S). Due to the importance of viral membrane fusion, S has been a popular target for developing vaccines and therapeutics. We discovered a simple peptide that inhibits infection by all major variants of SARS-CoV-2 with nanomolar efficacies. In marked contrast, widely used shorter peptides that lack a key N-terminal extension are about 100 x less potent than this peptide. Our results suggest that a simple peptide with a suitable sequence can be a potent and cost-effective therapeutic against COVID-19 and they provide new insights at the virus entry mechanism.

## Introduction

Viral fusion protein-derived peptides are potent entry inhibitors, as exemplified by enfuvirtide (T-20 or Fuzeon) (1, 2), an FDA-approved synthetic 36-amino-acid peptide targeting the human immunodeficiency virus type 1 (HIV-1) envelope glycoprotein 41, and more recent derivatives of this peptide (3). A similar strategy has been applied to target other viruses, including influenza (4), the original severe acute respiratory syndrome coronavirus (SARS-CoV) (5, 6), Middle East respiratory syndrome coronavirus (MERS-CoV) (7, 8), and the more recent severe acute respiratory syndrome coronavirus 2 (SARS-CoV-2) (9–11). In the case of SARS-CoV-2, peptides derived from the heptad repeat 2 (HR2) region of virus spike (S) protein show promising inhibition of viral entry (9–11). Following host receptor binding and proteolytic activation, trimeric S mediates virus entry by formation of the so-called HR1HR2 six-helix bundle that pulls together the host and virus membranes. HR2-based peptide inhibitors presumably promote bundle formation together with the heptad repeat 1 (HR1) region of S, precluding some or all of the intrinsic HR2 regions from binding to the HR1 regions and thus locking S into an extended, pre-hairpin fusion intermediate (12) that precludes viral entry.

To improve the efficacy of HR2-based inhibitors, various structural engineering approaches have been used. A SARS-CoV HR2-derived peptide, EK1, was mutated to enhance its stability, solubility, and antiviral activity (13), resulting in a half-maximal inhibitory concentration (IC_50_) of ~300 nM in a SARS-CoV-2 S based cell-cell fusion assay (9). Subsequently, a SARS-CoV-2 HR2 derived peptide was conjugated to polyethylene glycol and cholesterol, possibly to facilitate peptide targeting to the membrane, resulting in an IC_50_ of ~5 nM in an authentic SARS-CoV-2 infection assay (11). Additionally, a SARS-CoV-2 HR2 derived peptide was stabilized by the introduction of hydrocarbon staples, but the half maximal effective concentration values remain in the micromolar range (14). Notably, all three structural engineering approaches were limited to an identical 36 amino acid segment of SARS-CoV-2 S (residues 1168–1203).

Here, we report an extended, unmodified SARS-CoV-2 HR2 peptide (residues 1162–1203) with much improved properties relative to these previous attempts. During structural characterization of HR1HR2 bundles by high-resolution cryo-electron microscopy (cryo-EM), we observed the formation of a well-folded N-terminal region (residues 1159–1179) of HR2 in an extended conformation, outside the original range used for previous peptide studies. Based on this observation, we prepared an N-terminally extended HR2-derived peptide. Although this N-terminal extension confers only a modest increase in apparent binding affinity, the N-terminal extension much more substantially inhibits SARS-CoV-2 infection by ~100-fold with an IC_50_ value of ~1 nM as assessed by a cell-based fusion assay, suggesting that factors in addition to the thermodynamic stability of the bimolecular peptide:HR1 interaction determine the efficacy of these peptides in inhibiting the fusion process. The peptide also potently inhibits both VSV-SARS-CoV-2 chimera and authentic SARS-CoV-2 in the nanomolar range as determined by infection assays. Furthermore, the efficacy of the peptide is maintained for all major variants. The peptide has a very long inhibition lifetime (> 3 hours) after washout in virus infection assays, supporting the notion that multiple peptides bind to multiple pre-hairpin intermediate S proteins.

## Results

### An N-terminal extension of HR2 interacts with HR1 in a well-defined extended conformation

Previous high-resolution structures of the post-fusion HR1HR2 bundle of the Wuhan-Hu-1 strain of SARS-CoV-2 (GISAID accession ID: EPI_ISL_402124, here referred to as Wuh) revealed an overall six-helix bundle architecture in which the HR1 α-helices form a core with grooves into which the HR2 segments lie (PDB IDs 6lxt, 6m1v, 7rzq) (9, 15, 16). The cryo-EM structure of this HR1HR2 bundle (PDB ID 7rzq) (16) clarified several sidechain positions at high resolution and revealed that the N-terminal region of the HR2 fragment (residues 1164–1200) forms a well-defined extended conformation. Based on this observation, we further extended the N-terminal region, beginning at residue 1157, and determined the cryo-EM structure of this extended HR1HR2 complex (Fig. 1*B*, Fig. S1, and Table S1). Nearly all additional residues at the N-terminal region of HR2 are well ordered and form an extended conformation (Fig. 1*B*). We note that the densities for these N-terminal residues were not observable in the previous reconstruction (PDB ID 7rzq) that employed a shorter sequence at the N-terminus. The additional N-terminal residues of the HR2 fragment interact with the three HR1 α-helices. Moreover, the densities for residues 1164–1167 are also improved compared to the previous reconstruction. Residues V1164 and L1166 form hydrophobic interactions with the groove of the HR1 bundle (Fig. 1*B*). In addition, the backbone carbonyl oxygen and amide nitrogen atoms of residue G1167 form hydrogen bonds with the sidechain atoms of residue N969 of HR1 (Fig. 1*B*).

**Fig. 1.**
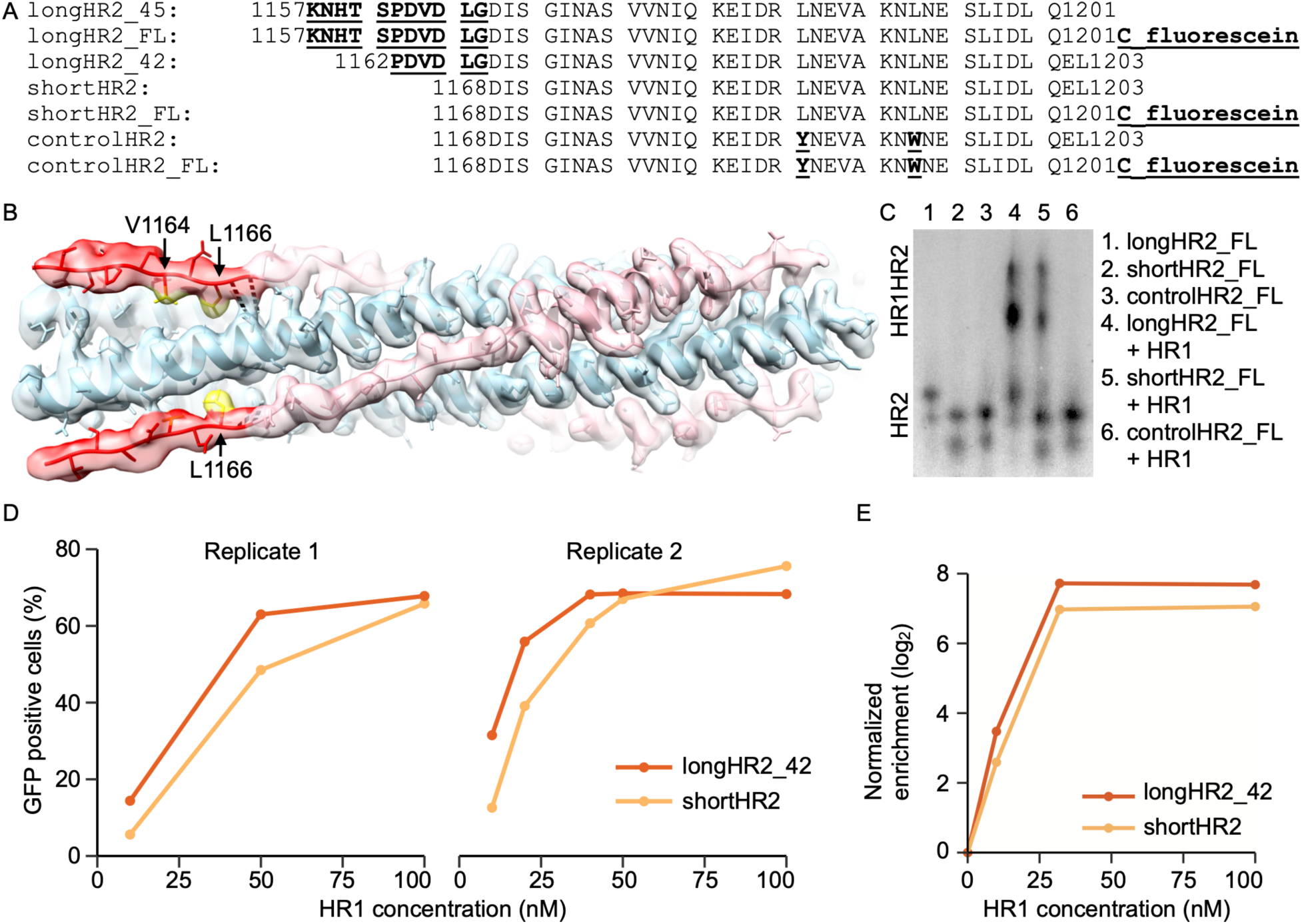
N-terminal extension of HR2 peptide interacts with HR1. (*A*) Sequences of all HR2 peptides used in this study (longHR2_45 corresponds to the Wuhan strain (GISAID Accession ID: EPI_ISL_402124). The numbers of residues, 45 and 42, are indicated in the naming for longHR2_45 and longHR2_42, respectively. The suffix “FL” indicates peptide labeling with fluorescein. Any difference from the traditionally used shortHR2 is indicated by underscore and bold font. (*B*) EM structure of longHR2_45 bound to HR1. Red: N-terminal extension of HR2. Pink: the rest of HR2. Blue: HR1. Yellow: hydrophobic sidechains of V1164 and L1166 interacting with HR1. Black dashed lines: hydrogen bonds between N969 in HR1 and G1167 in HR2. (*C*) CN-PAGE fluorescence imaging shows that longHR2_FL forms a more intense HR1HR2 bundle band than does shortHR2_FL, and that controlHR2_FL does not bind to HR1. (*D*) Bacterial surface display shows that longHR2 outcompetes shortHR2 for binding to HR1 at low HR1 concentrations. *E. coli* were incubated with different concentrations of GFP-tagged HR1 peptide (x axis) and then sorted using flow cytometry. The percentage of GFP-positive cells is indicated (y axis) for cells expressing longHR2 (red) or shortHR2 (orange). (***E***) mRNA display shows that longHR2_42 is more enriched than shortHR2 during affinity purification of HR1. mRNAs encoding longHR2_42 (red) or shortHR2 (orange) were translated in rabbit-reticulocyte lysate in the presence of different concentrations of HR1 (x axis), reverse transcribed, and sequenced using high-throughput sequencing. Resulting enrichments [log_2_(eluate / input)] were normalized to both a control peptide (with the sequence LKVLLYEEFKLLESLIMEILEYQKDSDIKENAEDTK, ref. (9)) and the no-HR1 control.

This cryo-EM structure suggests that the extended N-terminal region of HR2 confers additional stability to the complex. To test this, we introduced a cysteine in the C-terminus of HR2 to allow fluorescein-maleimide labelling and tracked the HR1HR2 bundle formation using clear-native polyacrylamide gel electrophoresis (CN-PAGE). We prepared two fluorescein-labeled HR2 peptides (Fig. 1*A*): longHR2_FL, corresponding to the longHR2_45 construct used in the present study, and shortHR2_FL, corresponding to the previously reported HR2 peptide (9, 11, 13, 14). As a control, we engineered controlHR2_FL that contains two point mutations in the helical region, L1186Y and L1193W, replacing these two leucine residues with bulky residues to promote clashes and disrupt interactions in the HR1HR2 bundle. Under identical experimental conditions, the longHR2_FL-containing HR1HR2 bundle forms a more intense high-molecular-weight band than the shortHR2_FL-containing HR1HR bundle, for which a larger pool of free HR2 remains in the low-molecular-weight region. As expected, the controlHR2_FL does not bind to HR1 (Fig. 1*C*). These results suggest that the N-terminal extension of HR2 indeed stabilizes the HR1HR2 bundle (Fig. 1*C*).

With an eye toward the eventual deployment of high-throughput screens for inhibitory peptides, we developed two assays to detect complex formation between HR2 variants and HR1. In the first assay, HR2 peptides were displayed on the surface of *E. coli* using an eCPX-protein scaffold. The cells were incubated with purified GFP-tagged HR1, and GFP-positive cells (which express HR2 peptides that bind HR1) were selected using fluorescence-activated cell sorting (17–19). The second assay employs mRNA display to detect complex formation (20, 21). mRNA display was used in an attempt to approximate co-translational assembly of the HR1HR2 bundle, and to facilitate future studies aimed at further optimization of the HR2 peptide. Synthetic mRNAs encoding HR2 or variants were prepared with a 3′ puromycin group, which promotes mRNA-peptide linkage during *in vitro* translation. The mRNAs were translated in the presence of hexahistidine-tagged HR1, and unbound peptides were separated from bound peptides by affinity purification with Ni-NTA beads. The selected mRNA-peptide fusions were reverse transcribed, PCR amplified, and sequenced using high-throughput sequencing. In order to compare across sequencing libraries, read counts from the selection were normalized to input counts to obtain enrichment values. These enrichments were then normalized both to a scrambled peptide (9) and to the background enrichment in a sample without HR1. Both assays show a modest (~2 fold) increase in apparent binding affinity for longHR2 compared to shortHR2, most pronounced at lower HR1 concentrations (Fig. 1 *D* and *E*, Fig. S2, and Table S2). These techniques provide the basis for further characterization and optimization of HR2-derived peptides.

### The N-terminal extension increases the inhibition activity 100-fold

In light of the observed interactions between HR1 and the N-terminal extension of HR2, we tested the effects of the N-terminal extension of HR2 on inhibiting the membrane-fusion function of SARS-CoV-2 S in a cell-cell fusion assay (22, 23). Two versions of N-terminally extended HR2 peptides, referred to as longHR2_45 and longHR2_42, inhibit cell-cell fusion with an IC_50_ of 1.6 nM and 1.3 nM, respectively, while the IC_50_ of the previously used HR2 peptide, shortHR2, is only 263.1 nM, consistent with a previous study (10) (Fig. 2). The controlHR2 peptide does not inhibit cell-cell fusion.

**Fig. 2.**
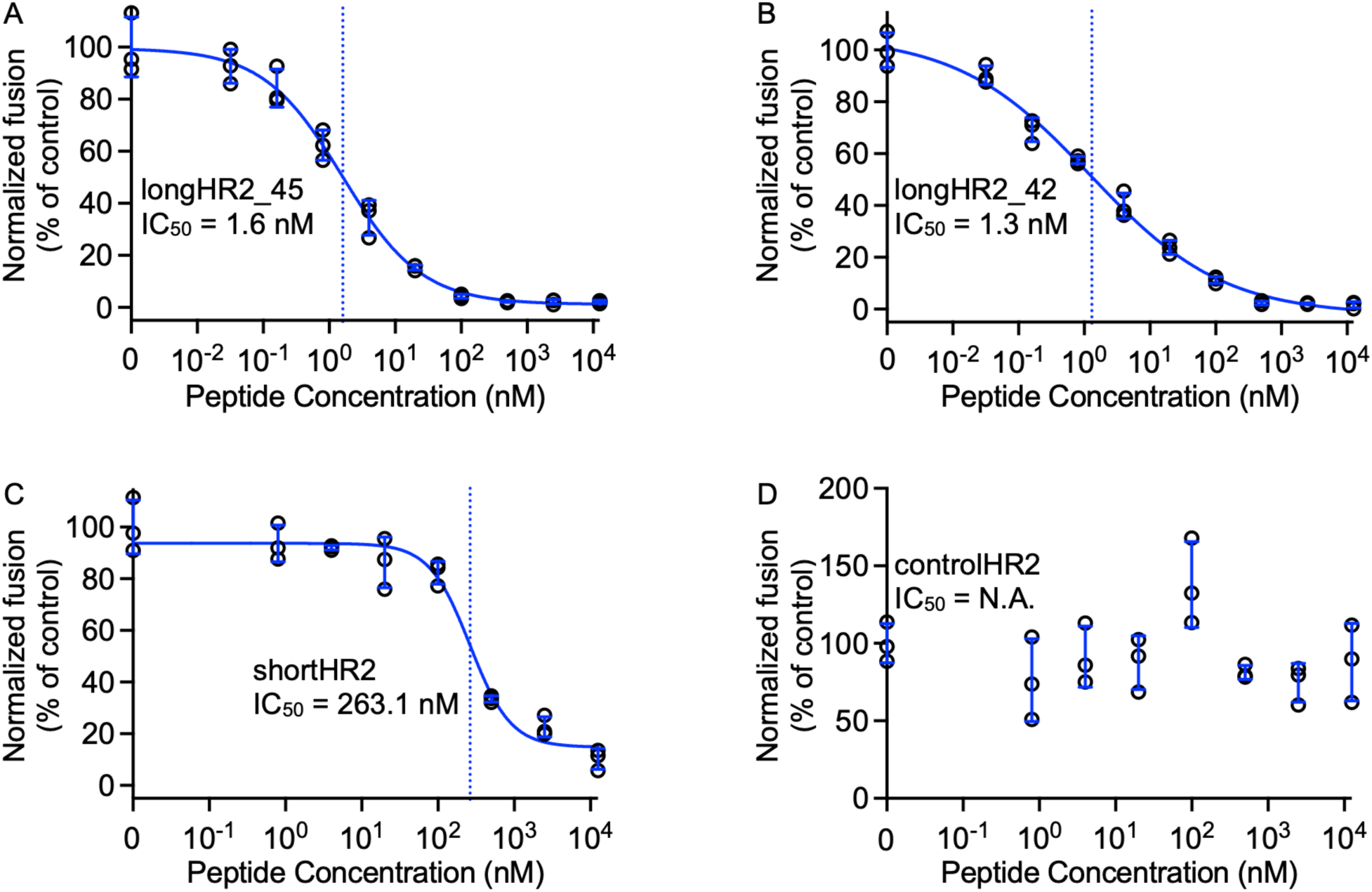
N-terminal extension of the HR2 peptide increases the efficacy by a hundred fold in the cell-cell fusion assay. Inhibitory activities of longHR2_45, longHR2_42, shortHR2, and controlHR2 in the cell-cell fusion assay are shown in panel *A, B, C*, and *D*, respectively. The raw data points are plotted as black circles, while the error bars, fitted curves, and vertical dash lines at IC_50_ are plotted in blue color (the same plotting scheme is used for all assays in Figs. 2–5).

As the C terminus of longHR2_42 is identical to that of the previously used shortHR2, any change in inhibition activity can only be attributed to the differences in the N-terminal region of HR2. We thus decided to use longHR2_42 for all subsequent experiments since it has a comparable IC_50_ value as longHR2_45 but with a slightly shorter sequence.

### Nanomolar inhibition in a VSV-SARS-CoV-2 chimera infection assay

The potential antiviral effect of the longHR2_42 peptide was explored using a chimeric VSV virus expressing a soluble eGFP infection reporter, where the glycoprotein of VSV was replaced with the S protein of SARS-CoV-2 with the sequence of the Wuh strain (VSV-SARS-CoV-2-Wuh). We note that this chimeric virus is able to replicate, unlike pseudotyped viruses. One round of VSV-SARS-CoV-2-Wuh infection was examined in VeroE6 cells overexpressing TMPRSS2 (VeroE6+TMPRSS2). Comparing longHR2_42 to shortHR2 we observed a decrease in IC_50_ to 1.1 nM (longHR2_42) from 198.5 nM (shortHR2) (Fig. 3). The inhibition of longHR2_42 for VSV-SARS-CoV-2 occurred across several cell lines with different expression levels of the host cell proteases cathepsin and TMPRSS2 necessary for S mediated infection. Specifically, Vero cells only possessing the cathepsin entry route, VeroTMPRSS2 cells containing both cathepsin and TMPRSS2 protease, and Calu-3 cells which are lung cells previously found to completely rely on the TMPRSS2 protease for VSV-eGFP-SARS-CoV-2 (24–26), were found to have complete inhibition of infection in the presence of 10 nM of longHR2_42 (Fig. S3). These results demonstrate the improved affinity of longHR2_42 and show this is independent of cell type or host cell proteases necessary for S protein cleavage.

**Fig. 3.**
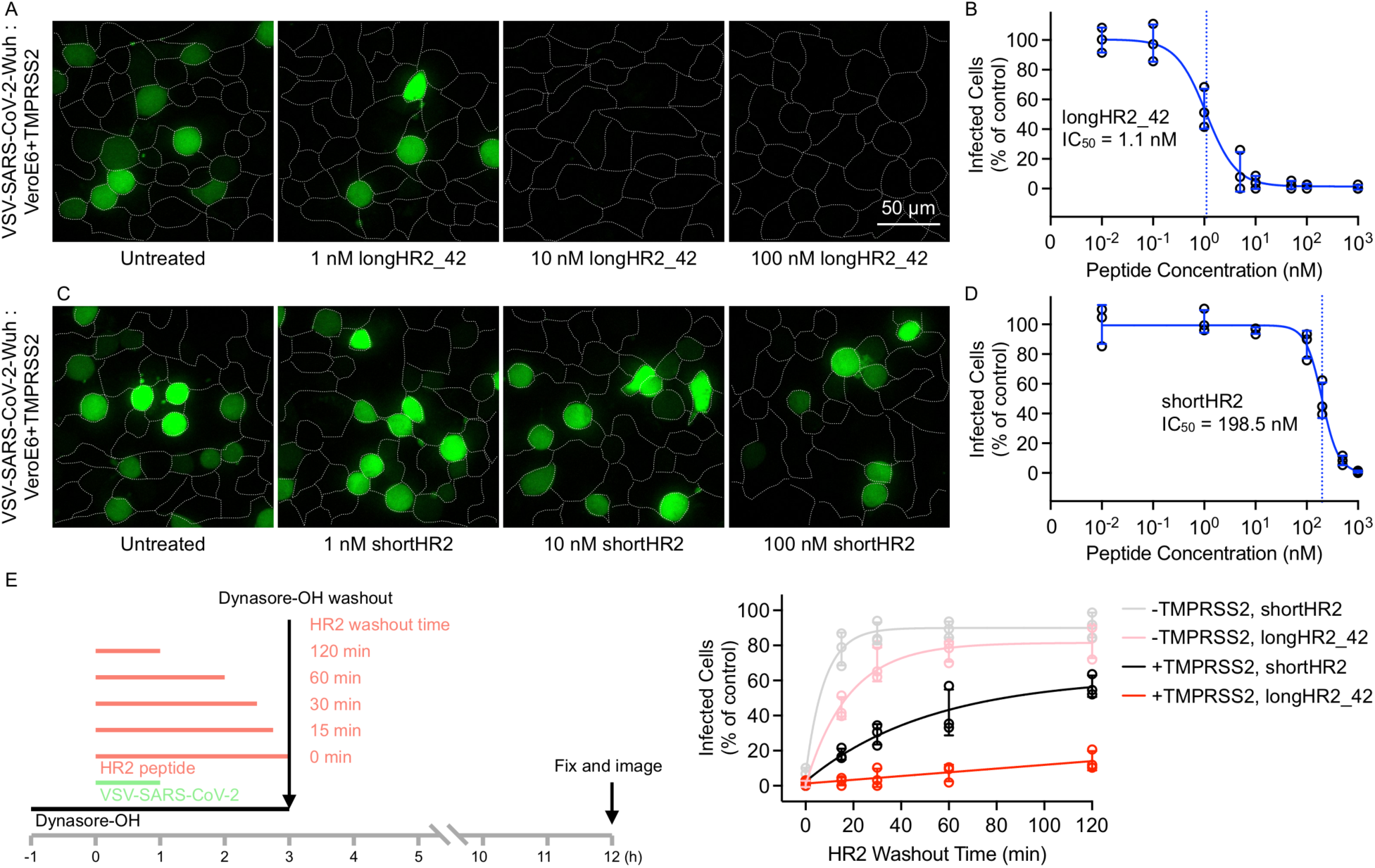
VSV-SARS-CoV-2 chimera infection assay confirms the strong inhibition by the N-terminally extended HR2 peptide. Inhibition of VSV-SARS-CoV-2-Wuhan infection (multiplicity of infection of 0.5) by the longHR2_42 (*A* and *B*) or shortHR2 (*C* and *D*) in VeroE6+TMPRSS2 cells. Virus and peptide were incubated with cells for 1 hour, washed, then fixed and imaged 8 hours of initiation of infection allowing for 1 round of infection to occur. Images are maximum intensity projections of 20 µm z-planes taken with 1 µm spacing using a spinning disc-confocal. Expression of a soluble eGFP (green) reported allowed for infected cells to be determined while cell outlines were obtained by from WGA-Alexa647 stain applied immediately prior to fixation (*A* and *C*). (*E*) Infection of VeroE6 and VeroE6+TMPRSS2 by VSV-SARS-CoV-2 (10 µg/mL viral RNA) where cells were inhibited with dynasore-OH for 4 hours, in which VSV-SARS-CoV-2 was added in the presence of 500 nM HR2 peptides and after 1 hour of binding the HR2 was washed out for different amounts of time. Schematic (left) and amount of infection (right) are shown.

### Binding of HR2 peptides to S is greatly enhanced in the presence of TMPRSS2

Dynamin-dependent endocytosis is required for SARS-CoV-2 infection independent of TMPRSS2 expression (27). Acute inhibition of dynamin by the chemical inhibitor dynasore-OH (28) allows for VSV-SARS-CoV-2 to bind to the cell surface; subsequent removal of dynasore-OH allows for infection to proceed. We allowed VSV-SARS-CoV-2 to bind to dynasore-OH treated VeroE6 or VeroE6+TMPRSS2 cells in the presence of 500 nM HR2 peptides for 1 hour. After 1 hour of binding, HR2 peptide was washed out for varying time intervals followed by washout of dynasore-OH (Fig. 3*E*). Examining the amount of infection 9 hours after the removal of dynamin inhibition allowed for inhibition lifetime after peptide washout to be observed (Fig. 3*E*). Incubation of cells with controlHR2 had no effect on infection as compared to the control while keeping dynasore-OH constant throughout the experiment completely inhibited infection (Fig. S4). In the absence of TMPRSS2, both peptides (shortHR2 and longHR2_42) are washed out within minutes. The expression of TMPRSS2 greatly increased the inhibition lifetime of both peptides after washout and made the binding of longHR2_42 effectively irreversible within our experimental conditions, exacerbating the difference between shortHR2 and longHR2_42. Trypsin treatment in TMPRSS2-negative VeroE6 cells showed similar inhibition as that of using TMPRSS2-expressing VeroE6 cells (Fig. S5), suggesting that either trypsin or TMPRSS2 can cleave the S protein in an appropriate way to trigger the transition of the S protein to the pre-hairpin intermediate.

### Nanomolar inhibition in an authentic SARS-CoV-2 infection assay

To validate these results using authentic SARS-CoV-2, we infected Caco-2 cells overexpressing the human ACE2 receptor (Caco-2+hACE2) with a patient isolate of the virus corresponding to the Wuh strain. Consistent with the results of the VSV-SARS-CoV-2 chimera infection assay, the IC_50_ value of longHR2_42 for the authentic SARS-CoV-2 infection assay at 8 h post infection is 1.5 nM (Fig. 4).

**Fig. 4.**
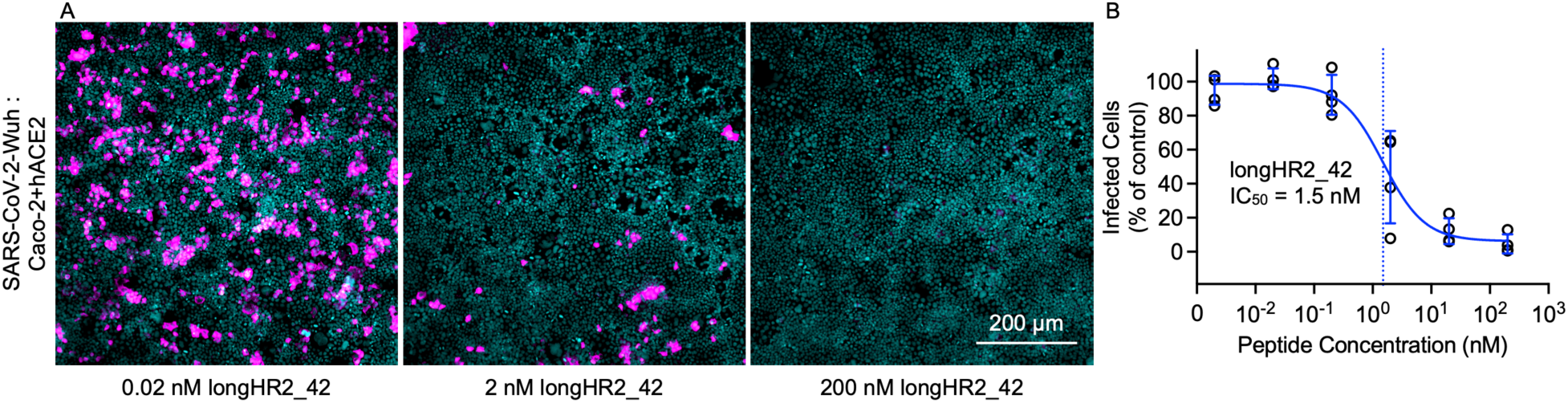
Authentic SARS-CoV-2 infection assay confirms the strong inhibition by the N-terminally extended HR2 peptide. Example images (*A*) and inhibition curve (*B*) from the infection of Caco-2+hACE2 cells by the Wuhan strain of SARS-CoV-2 in the presence of different concentrations of the longHR2_42 peptide. Nuclei are stained with Hoechst DNA dye (cyan), and infected cells are detected with an antibody specific for the viral N protein (magenta).

### Inhibition activity against all major SARS-CoV-2 variants to date

We next tested whether the longHR2_42 could inhibit infection by major SARS-CoV-2 variants using both the VSV-SARS-CoV-2 infection assay and the authentic SARS-CoV-2 infection assay. Patient isolates of Alpha, Delta, and Omicron variants were used to infect Caco2-hACE2 cells in the presence of increasing concentrations of longHR2_42 or controlHR2. The IC_50_ values remained at a similar single-digit nanomolar level in most cases (0.9 nM for both D614G and Delta in the VSV-SARS-CoV-2 infection assay, 0.6 nM for Alpha and 5 nM for Delta in the authentic SARS-CoV-2 infection assay) except for the Omicron variant, for which the IC_50_ was 4.1 nM in the VSV-SARS-CoV-2 infection assay and 15.6 nM in the authentic SARS-CoV-2 infection assay (Fig. 5, Figs. S6 and S7).

**Fig. 5.**
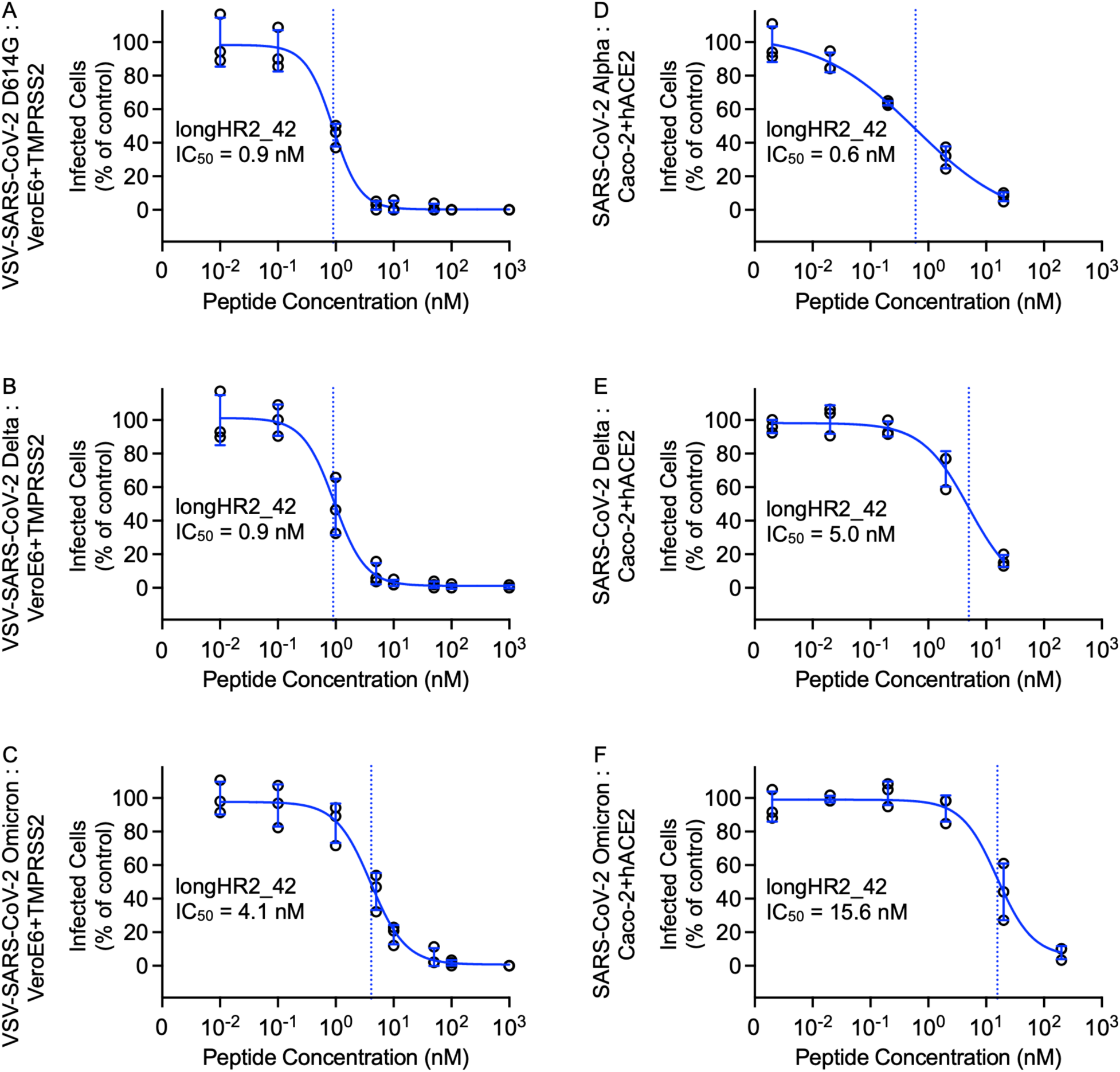
The N-terminally extended HR2 peptide inhibits infection by all major variants to date. (*A-C*) Inhibition of VSV-SARS-CoV-2-D614G (*A*), -Delta (*B*), or -Omicron (*C*) by longHR2_42 with a multiplicity of infection of 0.5 at 8 h post infection in Vero+TMPRSS2 cells. (*D-E*) Inhibition of SARS-CoV-2 Alpha (*D*), Delta (*E*), or Omicron (*F*) strain by longHR2_42 at 8 h post infection in Caco-2+hACE2 cells.

## Discussion

HR2-based fusion inhibitors represent an important class of viral therapeutics. Previous studies have focused on a short 36-amino-acid span of the HR2 region of S (9, 11, 14). Here, we report an N-terminally extended HR2 peptide, longHR2_42, that shows ~100-fold stronger inhibition than previously reported peptides without chemical modifications (9-11, 14). Specifically, we observed nanomolar inhibition by longHR2_42 in a cell-cell based fusion assay (Fig. 2), a VSV-SARS-CoV-2 chimera infection assay (Fig. 3), and an authentic SARS-CoV-2 infection assay (Fig. 4).

The N-terminal extension of HR2 interacts with HR1 in the postfusion HR1HR2 bundle in an extended conformation (Fig. 1*B*). However, the ~100-fold increased inhibitory efficacy of longHR2_42, compared to that of shortHR2, in fusion and infection assays, is not reflected in a corresponding increase in the apparent affinity of longHR2_42 for HR1, as measured by the three binding assays described here (Fig. 1 *C-E*). It is possible that the assays are not sensitive enough to detect changes in affinity when the affinity of shortHR2 for HR1 is already very strong. More likely, the discrepancy indicates that factors in addition to the bimolecular peptide:HR1 interaction determine the efficacy of these peptides in inhibiting the fusion process. Indeed, the SARS-CoV-2 S protein exists as a trimer, which cooperatively catalyze membrane fusion (Fig. 6). Single virus imaging of VSV-SARS-CoV-2 fusion indicated the simultaneous release of the S1 fragment from three to four S trimers followed by fusion of the virus with the host cell required slightly acidic media to trigger the required conformational changes in the S protein to the hairpin conformation (27). Thus, nine to twelve S proteins are probably necessary for fusion and a single inhibited protein of these nine to twelve proteins could therefore prevent fusion (Fig. 6). A small increase of the apparent binding affinity of the peptide would therefore lead to a large increase in potency and a large increase of the inhibition lifetime after peptide washout when the virus is attached to the surface of VeroE6+TMPRSS2 cells (Fig. 3*E*). Cleavage of the S protein by TMPRSS2 or cathepsin is a prerequisite for SARS-CoV-2 fusion (25). In the washout experiment (Fig. 3*E*) we initially prevent the action of cathepsin in the endosomal pathway by addition of the dynamin inhibitor dynasore-OH (28). The requirement of TMPRSS2 for the long inhibition lifetime after peptide washout suggests that cleavage by TMPRSS2 (or presumably also cathepsin) establishes a pre-hairpin intermediate of SARS-CoV-2 S that the peptide can readily bind to, in analogy to the pre-hairpin intermediate of HIV-1 gp-41 (12).

**Fig. 6.**
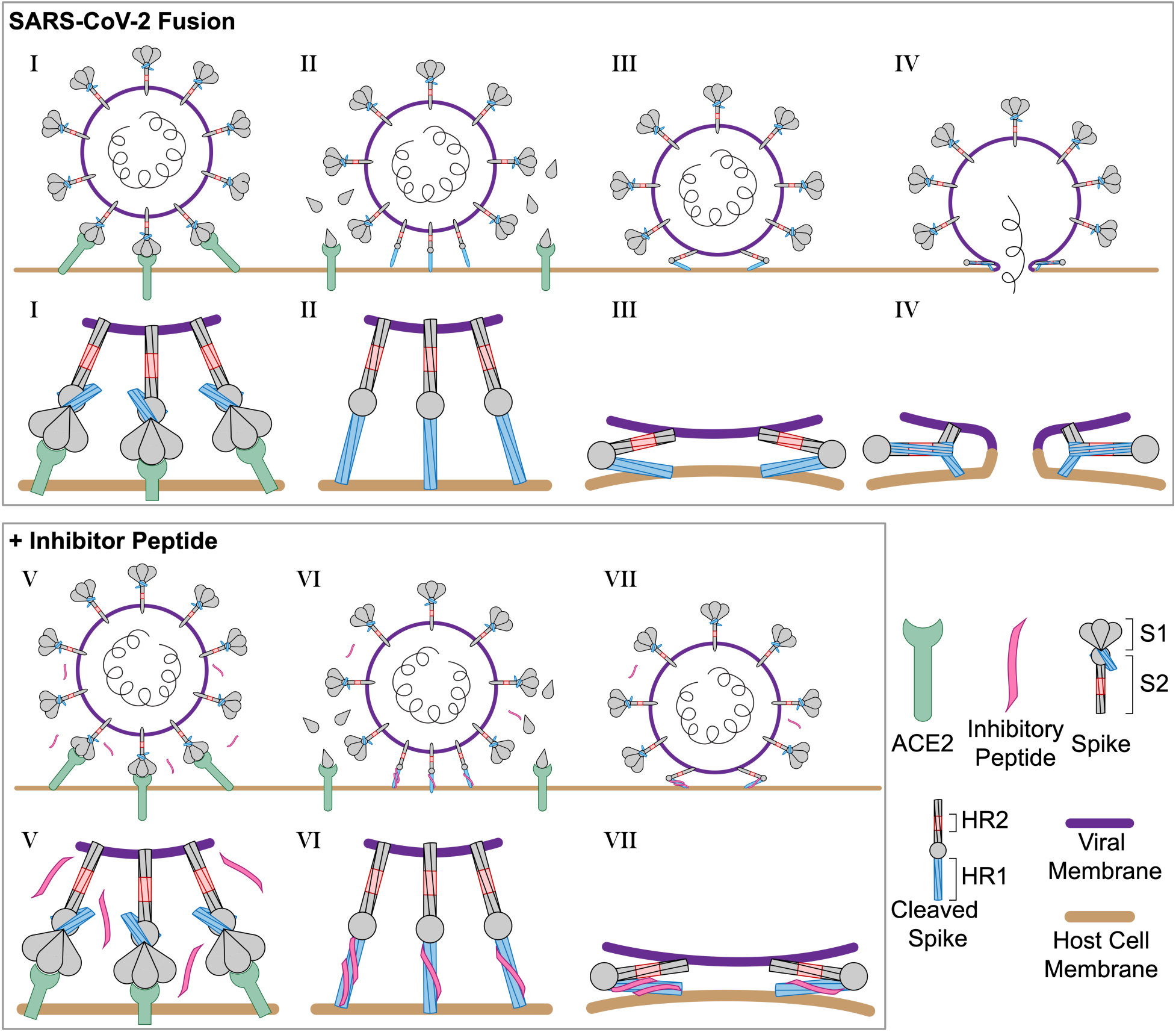
Schematic of SARS-CoV-2 infection and inhibition by HR2 peptides. (I) SARS-CoV-2 binds the host cell receptor ACE2 through an interaction with the S1 domain of the S protein. (II) After cleavage by host cell proteases the S1 domain is released and the S2 domain of the S protein extends into the host cell membrane. (III) Triggered by slightly acidic pH, the S2 domain folds back pulling the viral and host cell membrane into close proximity and (IV) the folding of the HR1 and HR2 domains catalyzes fusion of the viral membrane with the host cell membrane. (V) In the presence of the longHR2_42 inhibitor, the S protein engages the host cell receptor in the similar manner where (VI) after cleavage the longHR2_42 inhibitor binds the HR1 domain which (VII) prevents the folding of the HR1 and HR2 domains together and blocks membrane fusion. In this model, only a subset of the potential binding sites for the inhibitor need to be occupied in order to block fusion.

While only the TMPRSS2 protease cleavage occurs on the cell surface, the HR2 peptide inhibits infection in VeroE6 cells possessing only the cathepsin pathway (Fig. S3). It is unlikely that sufficient free peptide is encapsulated together with the virus to inhibit fusion throughout the endosomal pathway, but there may be some weak association of HR2 peptides with virus containing S in the prefusion state, leaving enough remaining peptide to inhibit the pre-hairpin intermediate after proteolytic activation. In support of this notion, the washout of HR2 peptide on the cell surface lacking TMPRSS2 shows that the peptide has an inhibition lifetime on the order of 15–20 minutes (Fig. 3*E*), suggestive of only a weak association between peptide and virus.

We also observed that longHR2_42 inhibits major SARS-CoV-2 variants (Fig. 5), although the activity is ~10-fold lower against the Omicron variant (Fig. 5). The D614G mutation present in all major variants is far from the HR1HR2 interface and no significant change is observed for the efficacy of the longHR2_42 against this mutant. Each of the Alpha and Delta variants has only a single mutation in the HR1HR2 bundle region (S982A and D950N, respectively) (Fig. S8). Neither mutation is involved in the interaction between HR1 and HR2 in the HR1HR2 bundle structures (9, 15, 16), and, as expected, there is no significant change in the efficacy of the longHR2_42 against these variants. In contrast, the Omicron variant has three mutations in the HR1 region—Q954H, N969K, and L981F—and all located at the interface between HR1 and the N-terminal extension of HR2 (Fig. S8), possibly explaining the somewhat weaker inhibition activity.

For the inhibition of the Delta variant, there exists an ~5-fold variation in the measured IC_50_ values between the VSV-SARS-CoV-2 infection assay and the authentic SARS-CoV-2 infection assay. This variation between different types of assays may be attributed to the fact that the mutated N protein from the Delta variant, which is absent in VSV-SARS-CoV-2 chimeric viruses, also significantly enhances infectivity (29). Moreover, the difference may also be due to the different cell lines used in the two assays.

In closing, we showed that an unmodified monomeric peptide sequence without any modifications or hydrocarbon staples shows nanomolar inhibition of SARS-CoV-2 infection by targeting the pre-hairpin intermediate of the S protein (Fig. 6). Additional potency could potentially be achieved by suitable extension of the peptide sequence. Further rounds of sequence optimization of longHR2_42 could improve the activity of the peptide and provide a platform for the development of new variant-specific peptides. Such designs could be important as antiviral therapeutics to combat the current SARS-CoV-2 pandemic as well as more distantly related viruses of concern.

## Materials and Methods

### Structure determination

The cryo-EM structure of the longHR2_45 bound to HR1 was determined following a molecular scaffolding method described previously (16). Briefly, the scaffolded HR1HR2 complex was generated by co-expressing the scaffolded HR1 and SUMO-tagged longHR2_45 in *E. coli* BL21(DE3) using auto-inducing LB medium (30), followed by Nickel affinity chromatography and size exclusion chromatography (SEC) with a Superose 6 Increase 10/300 GL column in 25 mM HEPES-Na, pH 7.4, 150 mM NaCl, 0.5 mM EDTA, 0.5 mM TCEP. The sample was concentrated to 20 µM, supplemented with 0.05% Nonidet P-40, and plunge frozen on a Quantifoil 2/1 holey carbon grid using a Vitrobot Mark IV (ThermoFisher). A total of 18,846 movies were recorded using a Titan Krios electron microscope (ThermoFisher) equipped with a K3 camera (Gatan) using the Serial-EM automation software (31), at a nominal magnification of 130,000 X and a pixel size of 0.3265 Å. Each movie contains 40 frames with a total electron dose of 47 e^-^/Å^2^. The data were processed using a combination of MotionCor2 (32), Gctf (33), EMAN2 (34), cryoSPARC (35), and RELION (36), as described previously (16). More details for data collection and processing are summarized in Fig. S1 and Table S1.

For model building, the PDB 7rzq was used as the template. The N-terminal extension of HR2 was first built in Coot (37) and then refined by the automated structure refinement protocol in Rosetta (38). The structure was then subjected to real space refinement (minimization_global, local_grid_search, adp) in PHENIX (39). Coot (37) was used for further fitting of sidechains and manual inspection.

### CN-PAGE and in-gel fluorescence imaging

The HR1 and HR2 peptides used for CN-PAGE were prepared in-house as described below. HR1 residues 910–988 of the Wuh strain SARS-CoV-2 S protein, and HR2 ranging from either residues 1157–1201 (longHR2_FL) or residues 1168–1201 (shortHR2_FL) of the Wuh strain SARS-CoV-2 S protein were cloned into the pETDuet-1 plasmid with N-terminal hexa-histidine and SUMO tags and a C-terminal cysteine for maleimide labeling. The peptides were recombinantly expressed in *E. coli* BL21(DE3) using auto-inducing LB medium (30). The HR1 peptide was purified by nickel affinity chromatography, SUMO protease cleavage, and SEC using a Superdex 30 HiLoad 16/60 column equilibrated with 20 mM sodium phosphate, pH 7.4, 150 mM NaCl, and 0.5 mM TCEP. The HR2 peptides were purified similarly but with an additional heat treatment at 95°C for 10 minutes following SUMO protease cleavage. The purified peptides were concentrated to ~1 mM, flash frozen in liquid Nitrogen and stored in a −80°C freezer for future use.

The maleimide labeling of the HR2 peptides was performed by mixing 100 µM peptide and 2.5 mM fluorescein-5-maleimide (ThermoFisher) in degassed buffer (50 mM sodium phosphate, pH 6.8, 150 mM NaCl, 0.05 mM TCEP), followed by overnight incubation at 4°C. Following labeling, free dye was removed by SEC using a Superdex 75 HR 10/300 GL column in PBS (137 mM NaCl, 2.7 mM KCl and 10 mM phosphate, pH 7.4) buffer. Labeled peptide was then further purified by high performance liquid chromatography (HPLC) with a C18 column in a gradient of water and acetonitrile supplemented with 0.1% trifluoroacetic acid followed by lyophilization and a final round of SEC using a Superdex 75 HR 10/300 GL column equilibrated with PBS buffer to remove any residual trifluoroacetic acid. The purified peptides were confirmed by mass spectrometry, concentrated to ~50 µM, protected from light, and stored at −80°C.

HR1HR2 complex formation was assessed by intrinsic fluorescence on CN-PAGE. Briefly, HR1 and HR2 peptides were diluted to 400 nM in PBS. 10 µL of HR1 was mixed with 10 µL of each HR2 peptide and incubated for 10 min at room temperature. The samples were then mixed with 5 µL of 4X native loading buffer (120 mM Tris pH 6.8, 20% glycerol, 0.02% bromophenol) and 5 µL of each sample were loaded onto an AnykD precast polyacrylamide gel (Bio-Rad). The gel was run in 25 mM Tris, 192 mM glycine, pH 8.3 at 70 V for 4 hours at 4°C. Protein LoBind tubes (Eppendorf) minimized potential adsorption of protein, and samples were protected from light throughout the entire process. The gel was then immediately imaged using an iBright 1500 imager (Invitrogen, excitation wavelength: 494 nm, emission wavelength: 512 nm).

### Purification of HR1 for the surface display and mRNA display assays

HR1 cDNA was inserted between a His_6_-SUMO sequence and the GFP gene and cloned into a pSMT3 vector. The SUMO-HR1-GFP fusion was expressed in BL21(DE3) cells by overnight induction with 0.5 mM isopropyl β-D-1-thiogalactopyranoside at 18°C. Cells were pelleted and then resuspended in a buffer containing 50 mM Tris, pH 8.0, 300 mM NaCl, 10 mM imidazole, 2 mM β-mercaptoethanol, and a cocktail of protease inhibitors. Cells were lysed by French press, and the lysates were clarified by centrifugation at 35,000 x g for 1 h. The SUMO-HR1-GFP fusion was purified using HisTrap^™^ Fast Flow column (Sigma). Eluted fractions were then dialyzed against a buffer containing 20 mM tris-HCl (pH 8.0), 150 mM NaCl, and 1 mM dithiothreitol at 4°C overnight. After dialysis, the sample was concentrated and gel-filtered in the same buffer on a HiLoad 16/60 Superdex 200 (GE Healthcare) column. Peak fractions that contain SUMO-HR1-GFP fusion were pooled and concentrated to 4 mg/ml through centrifugation (EMD Millipore).

### Surface display of the peptides

DNA encoding the peptides (longHR2_42, shortHR2, and the scrambled EK1 control), were cloned into a pBAD33 vector containing the enhanced circularly permuted OmpX protein (eCPX) (17). 100 ng of DNA was used to transform 100 µL of electrocompetent *E. coli* MC1061 cells. Details for the peptide expression can be found in ref. (19). After induction of peptide expression on the N terminus of the surface display scaffold eCPX, *E. coli* cells were resuspended in a buffer containing 50 mM HEPES, pH 7.5, 150 mM NaCl, and 0.2% BSA and SUMO-HR1-GFP at a concentration ranging from 0 to 100 nM. Cells were then washed and diluted 5-fold in the same buffer. Flow cytometry was used to detect and measure the binding of HR1 to the surface displayed peptides. The unlabeled bacterial cells were analyzed in Forward (FSC) versus Side-Scatter (SSC) mode as well as in fluorescence mode. Regions corresponding to cells were identified using FSC and SSC, and gates were set to include only these events in the following experiments with labeled samples. Gates for the binding analysis were defined using unlabeled cells (no HR1-GFP) and a scrambled peptide control. The distribution of GFP fluorescence (binding level) for the whole cell population was quantified (Fig. 1*D* and Fig. S2).

### mRNA display

To prepare DNAs for in vitro transcription, a 5′ constant sequence was attached to variants of interest using overlap-extension PCR. The constant sequence consisted of a hemagglutinin tag, a SUMO tag, and a FLAG tag and was amplified using primers that added a T7 RNA polymerase promoter and overlapped with the 5′ portion of the HR2 variants (forward: 5′ TCTAATACGACTCACTATAGGGACAATTACTATTTACAATTACAATGTACCCATACGATGTTCCAGATTACGCTGGCAGCAGCAGCAGCGGCCTGGTGCC 3′, reverse: 5′ CTTATCGTCGTCATCCTTGTAATCGGATCCACCAATCTGTTCTCTGTGA 3′). Products were column purified (Zymo DNA Clean and Concentrator, according to the manufacturer’s instructions). This amplicon (100 ng) was then included in three separate 50 µL PCRs of longHR2_42, shortHR2, and the scrambled EK1 control (9) contained within a library of single-residue substitutions (685 variants total, one substitution for each of the 19 amino acids at each of the 36 positions in shortHR2 and the WT sequence). These three additional PCRs were primed using the same forward primer that was used in creating the 5′ constant sequence and a reverse primer that added a sequence used for annealing the puromycin-linking primer (reverse: 5′ AATAGCCGGTGGGTTTTTGTTGTAGTCACCAG 3′), creating ~640 bp products that were agarose-gel purified.

DNA molecules were transcribed into mRNA using the Megascript T7 Transcription kit (ThermoFisher) according to the manufacturer’s instructions, using 3 pmol of DNA in a 60 µL transcription reaction for 4 h at 37°C. Samples were DNase treated (3 µL of Turbo RNase-free DNase, 2 U/µL, ThermoFisher) for 20 min at 37°C and then precipitated by addition of 290 µL of water, 39 µL of 3 M NaOAc, pH 5.5, 4 µL linear acrylamide (5 mg/mL, ThermoFisher), and 1 mL of ice-cold 96% ethanol followed by overnight incubation at −20°C.

To ligate the puromycin oligo, RNAs at ~10 µM were incubated with a 1.5-fold molar excess of each of two splint oligos (5′ TTTTTTTTTTTTAATAGCCGGTG 3′ and 5′ TTTTTTTTTTTTNAATAGCCGGTG 3′), a 2-fold molar excess of 3′ DNA adapter (5′ /5Phos/AAAAAAAAAAAAAAAAAAAAA/iSp9//iSp9/ /iSp9/AC C/3Puro/ 3′ where 5Phos denotes a 5′ phosphate, iSp9 denotes a triethylene glycol spacer, and 3Puro denotes the puromycin group, Integrated DNA Technologies), 6000 U of T4 DNA ligase (New England Biolabs), 200 U of Superase-In (ThermoFisher), and 1x T4 DNA ligase buffer (New England Biolabs) in a 300 µL reaction. RNA, splints, 3′ adapter, and buffer were first incubated at 50°C for 5 min and then cooled on ice for 5 min to anneal the splints. After addition of the additional reaction components, the full reaction was incubated at 37°C for 1 h, then brought up to 400 µL with water. Samples were phenol-chloroform extracted by addition of equal volume phenol:chloroform:isoamyl alcohol (25:24:1 v/v, ThermoFisher), centrifuged (10 min at 14,000 rpm at 4°C), washed with equal volume chloroform, centrifuged again, and precipitated. Following precipitation, ligated products (reaction efficiencies ~60%) were purified on a denaturing urea-PAGE gel containing 4% polyacrylamide.

Purified, puromycylated RNAs corresponding to longHR2_42, shortHR2, and the scrambled EK1 control were combined (~100 nM total final concentration) and translated in 50 µL reactions containing 12.5 µM supplemental tRNA (Promega), 50 U of Superase-In (ThermoFisher), 47.5 µL of nuclease-treated rabbit-reticulocyte lysate (Promega), and one of four concentrations of HR1 peptide (0, 10 nM, 32 nM, 100 nM). Reactions were incubated for 90 min at 30°C. KCl (500 mM final) and MgCl_2_ (60 mM final) were then added and the reaction was incubated for an additional 30 min at room temperature to promote peptide-RNA linkage.

To purify HR1-bound RNAs, translation reactions were mixed 1:1 with 2x Ni-NTA lysis/binding buffer (50 mM NaH_2_PO_4_, 300 mM NaCl, 10 mM imidazole, 0.05% Tween 20, pH 8.0), 10% v/v was removed as input, 20 µL of Ni-NTA magnetic agarose beads (Qiagen) were added, and reactions were incubated at 4°C with end-over-end rotation for 2 h. Following incubation, beads were immobilized via magnetization, supernatant was discarded, and beads were washed 3 times with 500 µL wash buffer (50 mM NaH_2_PO_4_, 300 mM NaCl, 20 mM imidazole, 0.05% Tween 20, pH 8.0). RNA–peptides were eluted in 50 µL of elution buffer (50 mM NaH_2_PO_4_, 300 mM NaCl, 250 mM imidazole, 0.05% Tween 20, pH 8.0), and eluates were reverse transcribed using Superscript III (ThermoFisher) using oligo-dT-20mer as the primer according to the manufacturer’s instructions.

Resultant cDNA was PCR-amplified using two rounds of PCR amplification, first with primers specific to the template (foward 5′ CACTCTTTCCCTACACGACGCTCTTCCGATCTNNNNCAAGGATGACGACGATAAG 3′ and reverse: 5′ TGACTGGAGTTCAGACGTGTGCTCTTCCGATCTNNNN GGTTTTTGTTGTAGTCACCAG 3′, where N denotes an equal mixture of the four nucleotides) and second with Illumina barcoding primers (forward: 5′ AATGATACGGCGACCACCGAGATCTACACXXXXXXXXACACTCTTTCCCTACACGAC 3′ and reverse: 5′ CAAGCAGAAGACGGCATACGAGATXXXXXXXXGTGACTGGAGTTCAGACGTG 3′, where XXXXXXXX denotes an 8 nt barcode). Products were purified using agarose-gel electrophoresis, quantified using PicoGreen (ThermoFisher), and sequenced on a MiSeq using a paired-end 300 cycle kit with V2 chemistry (Illumina).

Reads were assigned to each variant using Kallisto (40). The enrichment score for each variant was determined by the following formula: log_2_(Eluate_count_Variant_ / Input_count_Variant_) − log_2_(Eluate_count_scrambled_ / Input_count_scrambled_) − log_2_(Eluate_count_0nM_HR1_ / Input_count_0nM_HR1_).

### Peptide synthesis and characterization

All peptides used for the subsequent assays were synthesized by GenScript USA Inc. The HPLC and liquid chromatography mass spectrometry (LC-MS) profiles provided by the manufacturer for longHR2_45, longHR2_42, shortHR2, and controlHR2 are shown in Figs. S9–S12, respectively.

In addition, we performed SEC and size exclusion chromatography coupled with multi-angle light scattering (SEC-MALS) to further characterize the peptides in the PBS buffer. Peptide powder was first dissolved in dimethyl sulfoxide (DMSO) to ~5 mg/mL. Subsequently, DMSO was exchanged to PBS buffer by three rounds of dilution and concentration. The dilution factor was ~15 for each round and the centrifugal concentrator (Merck Millipore Ltd.) had a molecular weight cut-off at 3 kDa. SEC and SEC-MALS were performed in PBS buffer using a Superdex 75 10 300 GL column and a wtc-010S5 column (Wyatt Technology Corporation), respectively. The SEC and SEC-MALS profiles for longHR2_45, longHR2_42, shortHR2, and controlHR2 are shown in Figs. S9–S12, respectively.

The concentration of peptide stock solution was determined by absorption measurement at 205 nm using a Nanodrop instrument (ThermoFisher), and confirmed using a Pierce(tm) BCA protein assay kit (ThermoFisher) and a Qubit protein assay kit (ThermoFisher).

### HEK cell-cell fusion assay

We optimized the cell-cell fusion assay (22, 23) based on the α-complementation of *E. coli* β-galactosidase for comparing the inhibitory efficacy of different peptides with higher throughput. Suspension culture Expi293F cells (ThermoFisher) were grown to a density of 1~2×10^6^ cells/mL in FreeStyle 293 expression medium (ThermoFisher) supplemented with 0.1 mg/mL penicillin-streptomycin antibiotics. The cells were then pelleted, resuspended in medium without antibiotics to a density of 1×10^6^ cells/mL, and allowed to recover at 37°C for 30 min. One group of cells was then co-transfected using polyethyleneimine (PEI, Sigma) (125 µg PEI / mL cells) with Wuh strain full-length SARS-CoV2 S protein construct (12.5 µg DNA / mL cells) and the α-fragment of *E. coli* β-galactosidase construct (12.5 µg DNA / mL cells) to generate the S protein expressing cells. Using the same amount of PEI, the other group of cells was co-transfected with the full-length ACE2 (12.5 µg DNA / mL cells) construct and the ω-fragment of *E. coli* β-galactosidase construct (12.5 µg DNA / mL cells) to generate the ACE2 receptor expressing cells. As a negative control, two additional groups of cells were transfected with either the α-fragment or the ω-fragment of *E. coli* β-galactosidase construct alone. After incubation at 37°C for 24 hr, the cells were pelleted. The S-expressing cells were resuspended FreeStyle 293 expression medium supplemented with different concentrations of peptide (50 µL, 2×10^6^ cells/mL), respectively. The ACE2-expressing cells and negative control cells were resuspended in 50 µL FreeStyle 293 expression medium to be 2×10^6^ cells/mL. S-expressing and ACE2 cells or α-fragment and ω-fragment cells were then mixed in a 96-well plate (Greiner bio-one) to initiate cell-cell fusion at 37°C for 2 hr. Fusion was arrested by adding 100 µL β-galactosidase substrate from the Gal-Screen reporter system (Invitrogen). The mixture was incubated at 37°C in the dark for 1 h before recording luminescence using a Tecan Infinite M1000.

### Purification of VSV-SARS-CoV-2 chimeras

Recombinant VSV chimera with glycoprotein G replaced with the SARS-CoV-2 S protein with the sequence of the Wuhan-Hu-1 strain, the D614G mutation in the Wuhan-Hu-1 strain, the Delta strain, or Omicron strain of SARS-CoV-2 (VSV-SARS-CoV-2) and expressing a soluble eGFP infection reporter was generated as described previously (24, 27, 41). VSV-SARS-CoV-2 was grown by infecting 12–18 150 mm dishes of MA104 cells at an MOI of 0.01. Supernatant was collected at 48 h post infection. The supernatant was clarified by low-speed centrifugation at 1,000 x g for 10 min at 4°C. Virus and extracellular particles were pelleted by centrifugation in a Ti45 fixed-angle rotor at 30,000 x g) for 2 h at 4°C. The pellet was resuspended in NTE buffer (100 mM NaCl, 10 mM Tris-HCl pH 7.4, 1 mM EDTA) at 4°C. The resuspended pellet was layered on top of a 15% sucrose-NTE solution and pelleted in an SW55 swinging-bucket rotor at 110,000 x g for 2 hours at 4°C. The virus was resuspended in NTE overnight at 4°C, then separated on a 15–45% sucrose-NTE gradient by ultracentrifugation in an SW55 swinging-bucket rotor at 150,000 x g for 1.5 h at 4°C. The predominant band observed in the lower one-third of the gradient was then extracted by side puncture of the gradient tube. Virus was then diluted in NTE and pelleted by ultracentrifugation in a Ti60 fixed-angle rotor at 115,000 x g for 2 h at 4°C. The VSV-SARS-CoV-2 pellet was resuspended overnight in NTE in a volume of 0.5 mL and stored at 4°C for subsequent experiments.

### VSV-SARS-CoV-2 infection assay

Glass slides (18 mm) were cleaned, mounted with 3-mm polydimethylsiloxane (PDMS) wells, and sterilized as previously described (24). On the day prior to the experiment, VeroE6 cells overexpressing TMPRSS2 (Vero+TMPRSS2) were plated in PDMS wells on class slide and stored in a 6 well plate at a density to achieve 70–80% confluence on the day of the experiment. On the day of experiments medium was removed, virus was diluted into media containing the desired concentration of indicated peptide at a final VSV-SARS-CoV-2 concentration of 0.5 µg/mL viral RNA (an MOI of ~0.5 infectious units (IFU) in Vero+TMPRSS2 cells), and then immediately added to the desired PDMS well in a volume of 10 µL. Medium was left in 5-well plate outside of the PDMS well at a level lower than the height of the PDMS well to maintain humidity and prevent evaporation. VSV-SARS-CoV-2 was incubated with the cells for 1 hr, then the cells were washed twice with medium to remove both unbound virus and inhibitor, and the well was then filled with fresh medium. In all experiments, cells were kept at 37^°^C with 10% CO_2_, and the medium was prewarmed to 37°C. At 8 h post infection the medium was removed; cells were stained with 5 µg/mL WGA-Alexa647 in PBS for 30 s at room temperature. Cells were then washed twice with PBS, fixed with 4% paraformaldehyde in PBS for 15 min, and then washed three times with PBS. Infected cells were imaged using a spinning-disk confocal microscope with a 40x oil objective and a pixel size of 0.33 µm where 20 optical planes were taken at 1 µm apart for every field of view (42). Cells were considered infected when they displayed a cytosolic eGFP fluorescence signal with a relative intensity 1.4 times that of the background of uninfected cells. Example images are maximum-intensity projections of the cell volume where the cell outline was obtained by tracing the WGA-Alexa647 membrane label.

### HR2 washout of dynamin inhibited VSV-SARS-CoV-2 infection

VeroE6 or VeroE6+TMPRSS2 cells were incubated with 40 µM dynasore-OH for 1 hour in serum free DMEM media containing 25mM HEPES, pH 7.2. Subsequently, VSV-SARS-CoV-2 (10 µg/mL of viral RNA) was added to the cell in the presence of 500 nM of peptides while keeping the concentration of dynasore-OH constant. After one hour, the virus was washed from all cells, and the HR2 peptides were washed out by exchanging the media 2 times with serum free DMEM containing 40 µM dynasore-OH (25mM HEPES, pH 7.4) when washing the virus (2 hour washout), an hour after removing the virus (1 hour washout), 1.5 hours after removing the virus (30 min washout), 1.75 hours after removing the virus (15 min washout) and 2 hours after removing the virus (0 min washout) (Fig. 3*E*). After completing the last HR2 washout in the series, dynasore-OH was removed by exchanging the serum free DMEM (25mM HEPES, pH 7.4) media 2 times. Virus and HR2 peptides were delivered to cells in 10 µL volumes to PDMS wells on a cover glass in a 6 well plate. When washes of the HR2 were performed the entire 6 well plate was filled with 3 mL of DMEM media (keeping dynasore-OH constant) followed by removal and replacement of media. The large excess volume of 3 mL compared to that of the PDMS well of 10 µL should have effectively removed and diluted any excess HR2 peptide. At 9 hours after dynasore-OH washout, the cells were stained with WGA-Alexa647 and fixed with 4% paraformaldehyde in PBS. Infection was then determined by imaging cells for GFP signal on a spinning-disc confocal as done with other VSV-SARS-CoV-2 infection assays. The infection was normalized to a control experiment of dynasore-OH washout in the absence of HR2 peptide inhibition. Trypsin treatment for the HR2 washout experiment (Fig. S5) was performed using the same procedure as previously published (27).

### SARS-CoV-2 infection assay

The SARS-CoV-2 infection assay was performed as described previously (24). Briefly, all experiments with SARS-CoV-2 were performed in biosafety level 3 (BSL3) facilities at the University of Helsinki with appropriate institutional permits. Nasopharyngeal samples were obtained under Helsinki University Hospital laboratory research permit 30 HUS/32/2018§16. Viruses were isolated and amplified once in VeroE6+TMPRSS2 cells for 48 h in MEM containing 2% FCS, 2 mM glutamine, and 1% penicillin/streptomycin, and viruses stored at −80 C. Virus titers were determined by a plaque assay in VeroE6+TMPRSS2 cells and the genome sequence of all virus stocks confirmed by deep sequencing (43). Caco2-hACE2 cells (44) were maintained in DMEM, supplemented with 10% FBS, 2 mM l-glutamine, 1% penicillin-streptomycin, and seeded 48 h before treatment at a density of 15,000 cells per well in 96-well imaging plates (catalog number 6005182; PerkinElmer). Inhibitors, or the DMSO control, were added either 15 min before infection of Caco-2-ACE2 cells at an MOI of 0.5 IFU per cell (corresponding to MOI 5 in VERO-E6 TMPRRSS2 cells). Infections were carried out for 8 h at 37°C with 5% CO_2_. Cells were then fixed with 4% paraformaldehyde in PBS for 30 min at room temperature, and the plates UV sterilized before removal from the BSL3 facility and processing for immunodetection of viral N protein, automated fluorescence imaging, and image analysis. Briefly, viral NP was detected with an in-house-developed rabbit polyclonal antibody (34), counterstained with Alexa Fluor 647-conjugated goat anti-rabbit secondary antibody, and nuclear staining was done using Hoechst DNA dye. Automated fluorescence imaging was done using a Molecular Devices Image-Xpress Nano high-content epifluorescence microscope equipped with a 10× objective and a 4.7-megapixel CMOS (complementary metal oxide semiconductor) camera (pixel size, 0.332 µm). Image analysis was performed with CellProfiler-3 software (www.cellprofiler.org). Automated detection of nuclei was performed using the Otsu algorithm inbuilt in the software. To automatically identify infected cells, an area surrounding each nucleus (5-pixel expansion of the nuclear area) was used to estimate the mean fluorescence intensity of the viral NP immunolabeled protein in each cell, using an intensity threshold such that <0.01% of positive cells were detected in noninfected wells.

### Statistics and Data Analysis

Data from the HEK cell-cell fusion assay, VSV-SARS-CoV-2 infection assay, and authentic SARS-CoV-2 infection assay contain three independent replicates at each inhibitor concentration. For the HEK cell-cell fusion assay, the normalized fusion was calculated as (Luminescence_(+inhibitor)_ - Luminescence_(α&ω)_) / (Luminescence_(+PBS)_ - Luminescence_(α&ω)_), where “+inhibitor” or “+PBS” refers to adding inhibitor or PBS to the mixture of the cells expressing α and S, and the cells expressing ω and ACE2, and “α&ω” refers to the mixture of the cells expressing α only and the cells expressing ω only. The counted infected cells were normalized by that of controlHR2 in the VSV-SARS-CoV-2 infection assay, and by that of shortHR2 at the lowest concentration (2 pM) in the authentic SARS-CoV-2 infection assay. After the normalization, the arithmetic means of the three replicates were used to fit the inhibition curves and estimate the IC50 values using the nonlinear regression of inhibitor concentration vs. response in GraphPad Prism version 9.1.0 for macOS (GraphPad Software, San Diego, California USA, www.graphpad.com). The fitting model is Y = Bottom + (Top - Bottom) / (1 + (IC50 / X) ^ HillSlope), where Y is the inhibition, X is the inhibitor concentration, Bottom is the maximal inhibition, and Top is the minimal inhibition.

### Figure preparation

The figures of PDB structures and maps were made in UCSF Chimera (45). The data fitting of all inhibition assays was performed and plotted using GraphPad Prism version 9.1.0 for macOS (GraphPad Software, San Diego, California USA, www.graphpad.com). Analysis and plots for the binding assays were performed using R (www.r-project.org).

## Data availability

The EM map and model of the HR1HR2 complex have been deposited in the EMDB (EMD-27098) and PDB (PDB ID: 8czi), respectively. Raw and processed RNA-seq data (used for mRNA display) is available at the GEO, accession number GSE203229.

## Acknowledgments

We thank Sondra Schlessinger for stimulating discussions, Bing Chen for kindly providing plasmids and protocols for the cell-cell fusion assay, the Stanford cryo-Electron Microscopy center support team, particularly Elizabeth Montabana and Chensong Zhang, for help with data collection during the COVID-19 pandemic, and staff at HUSLAB Virology and Immunology for providing samples for virus isolation, as well as support by the Academy of Finland 335527 / 336490 and HUH Funds TYH2021343 to G.B. and O.V., the Harvard Virology Program NIH training grant T32 AI07245 postdoctoral fellowship to A.J.B.K., the HHMI Helen Hay Whitney Foundation fellowship to K.I.W., the Damon Runyon fellowship DRG-2429-21 to T.J.E., NIH Maximizing Investigators’ Research Award (MIRA) GM130386 to T.K., NIH Grant AI163019 to T.K. (and Sean PJ Whelan), funds from Danish Technical University to T.K., and SANA to T.K.. This article is subject to HHMI’s Open Access to Publications policy. HHMI lab heads have previously granted a nonexclusive CC BY 4.0 license to the public and a sublicensable license to HHMI in their research articles. Pursuant to those licenses, the author-accepted manuscript of this article can be made freely available under a CC BY 4.0 license immediately upon publication.

## Figures and Legends

**Fig. S1.**
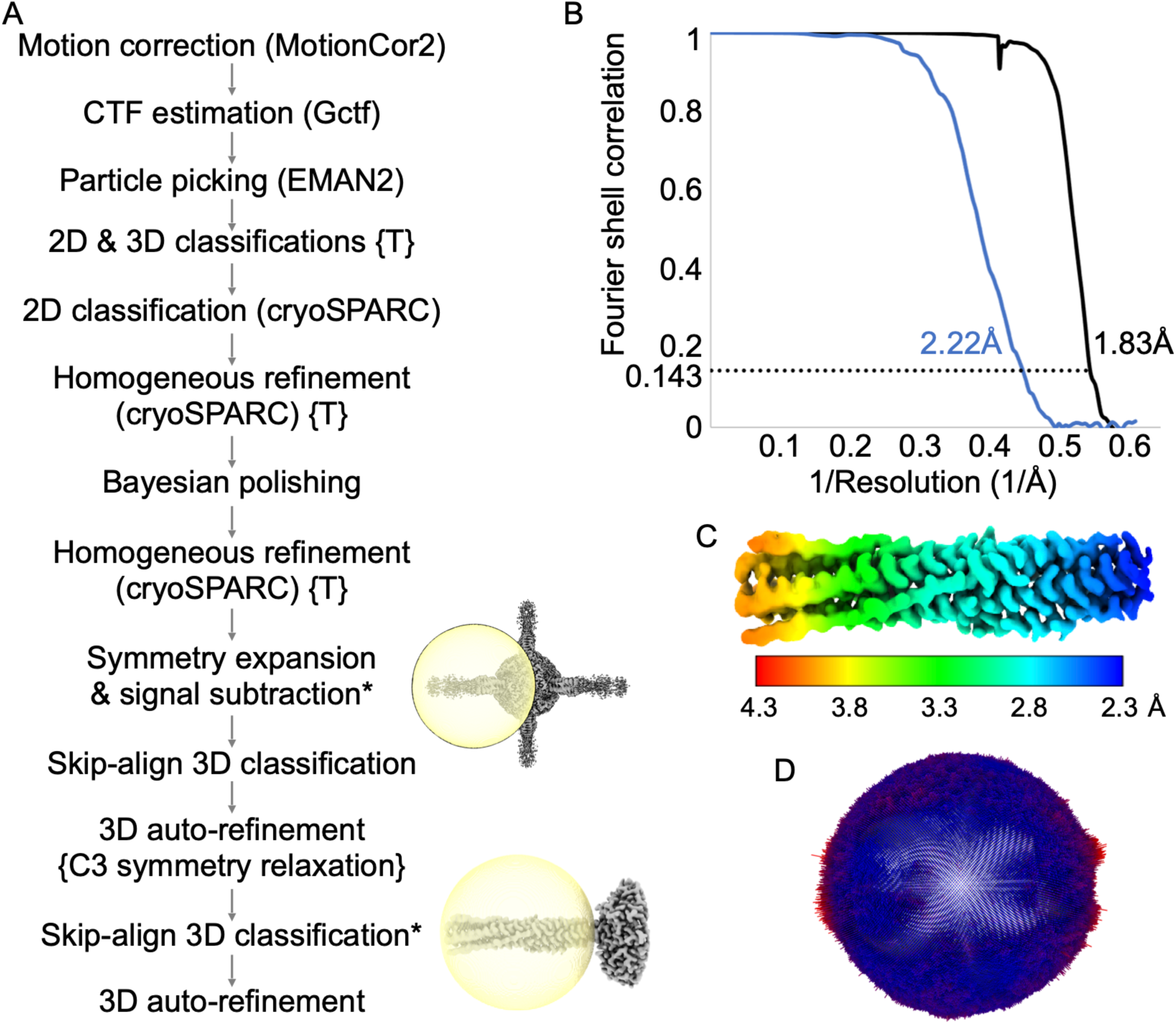
Cryo-EM structure determination. (*A*) Workflow of cryo-EM data processing. RELION was used for each step unless another program is indicated in parenthesis. Symmetry was imposed or relaxed as indicated in curly brackets. The steps that used a manually generated mask (rather than the default spherical mask in RELION or the default dynamic masks in cryoSPARC) are indicated with a star sign and an image showing the manually generated mask (yellow) and an average map (gray). (*B*) Fourier shell correlations (FSC) of the final refinement step with cryoSPARC (second “homogeneous refinement” using cryoSPARC, black curve) and the final local refinement with RELION (last 3D auto-refinement, blue curve. Note that these FSC calculations were performed using the default spherical mask that covers both the HR1HR2 bundle and part of the scaffold. (*C*) The final reconstruction of the HR1HR2 bundle colored by local resolutions. (*D*) Distribution of the particles’ orientations in the final reconstruction, depicted in the same orientation as the map in **c**. The length of each bar is proportional to the number of particles oriented in that direction. The bars are also colored based on the length, with red meaning more particles and blue meaning less particles.

**Fig. S2.**
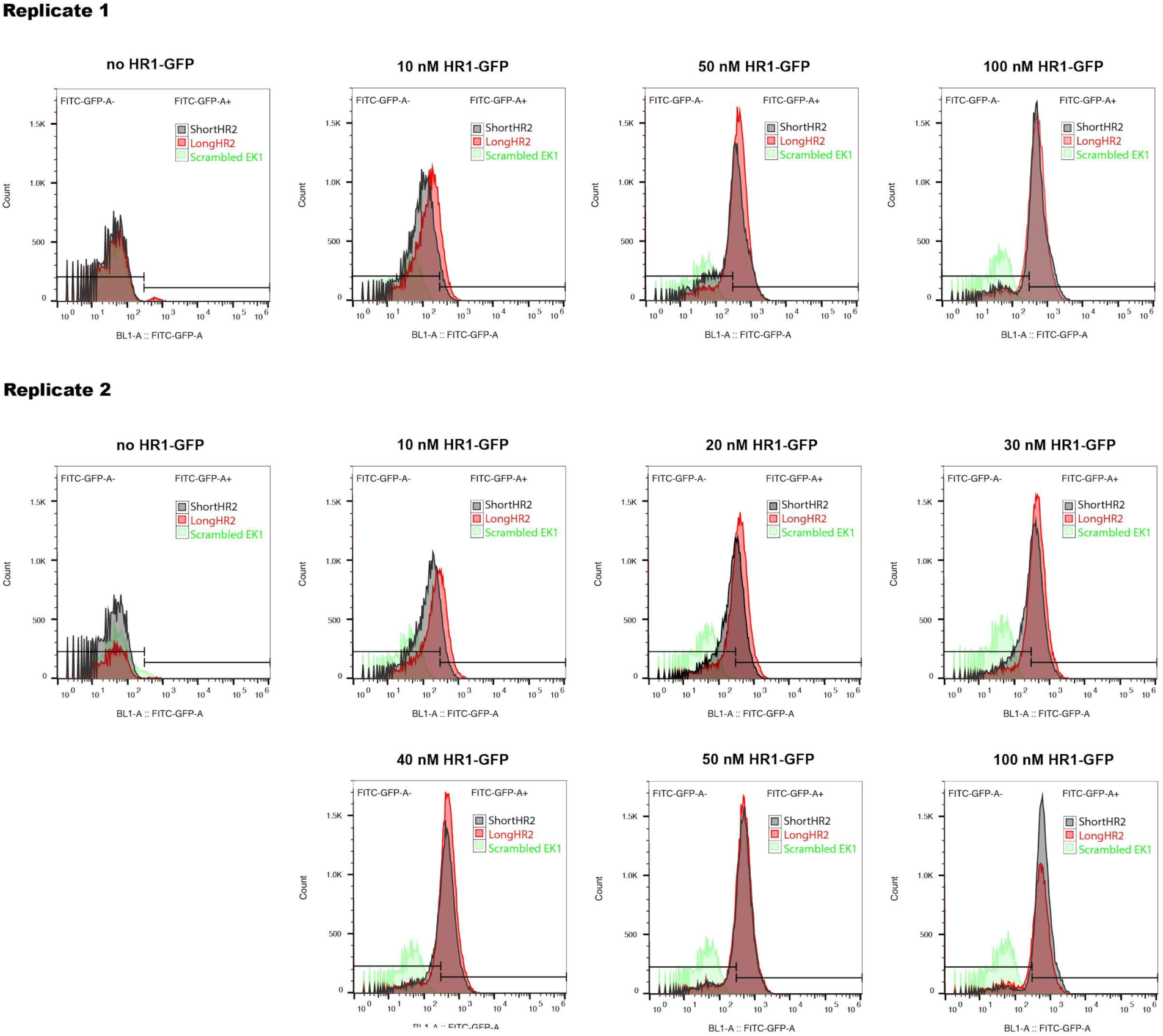
Flow cytometry histograms of HR1-GFP binding by surface-displayed longHR2_42, shortHR2, and the scrambled EK1 control. Cells displaying either peptide were incubated with different concentrations of HR1-GFP protein and the GFP signal was quantified to compare the binding levels. Regions corresponding to cells were identified using Forward (FSC) and Side-Scatters (SSC), and gates were set to include only these events in the experiments with labeled samples. Gates for the binding analysis were defined using unlabeled cells (no HR1-GFP) and the scrambled EK1 control.

**Fig. S3.**
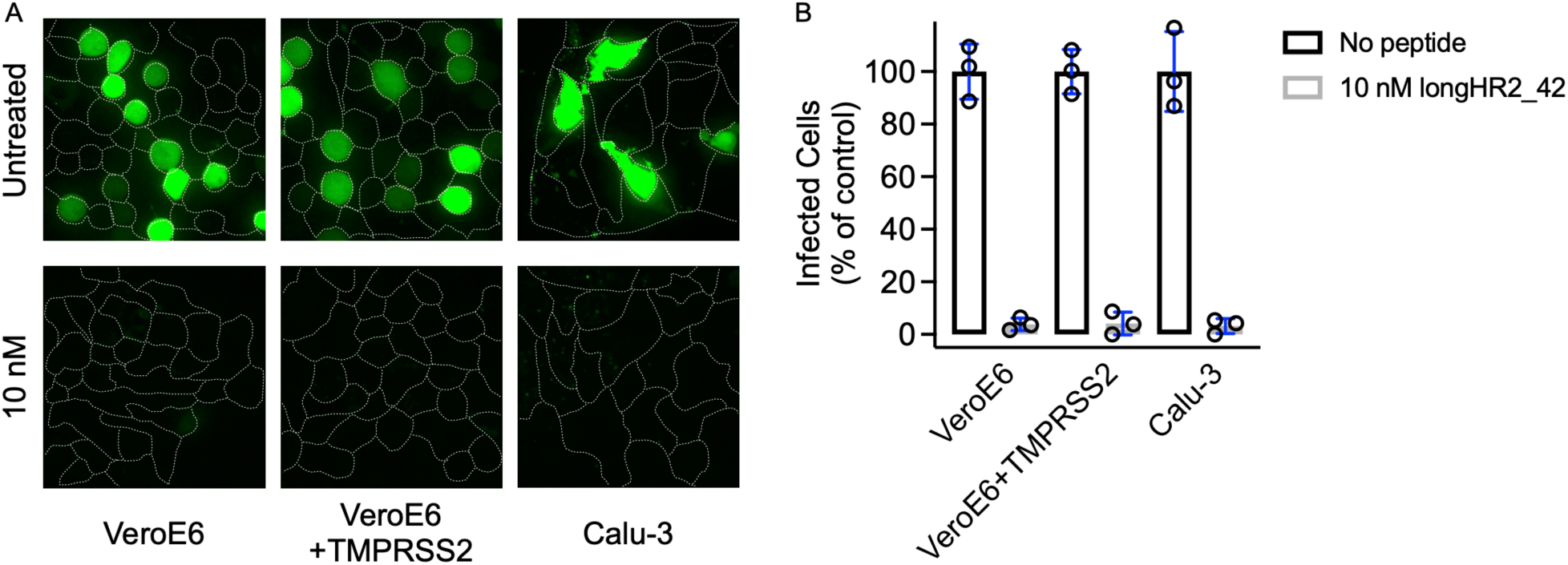
The strong inhibitory effect of longHR2_42 on VSV-SARS-CoV-2 is independent of cell types with differential expression of host cell proteases (*A* and *B*). Infection (MOI of 0.5) was determined in the absence or presence of 10 nM longHR2 in VeroE6 (cathepsin only), VeroE6+TMPRSS2 (cathepsin and TMPRSS2), and the human lung cell line Calu-3 (TMPRSS2 only). (*A*) Example images are maximum intensity projections of 20 µm with 1 µm z-plane spacing taken on a spinning disc-confocal with eGFP expression allowing for infection to be determined. Cell outlines were obtained from a WGA-Alexa647 membrane stain.

**Fig. S4.**
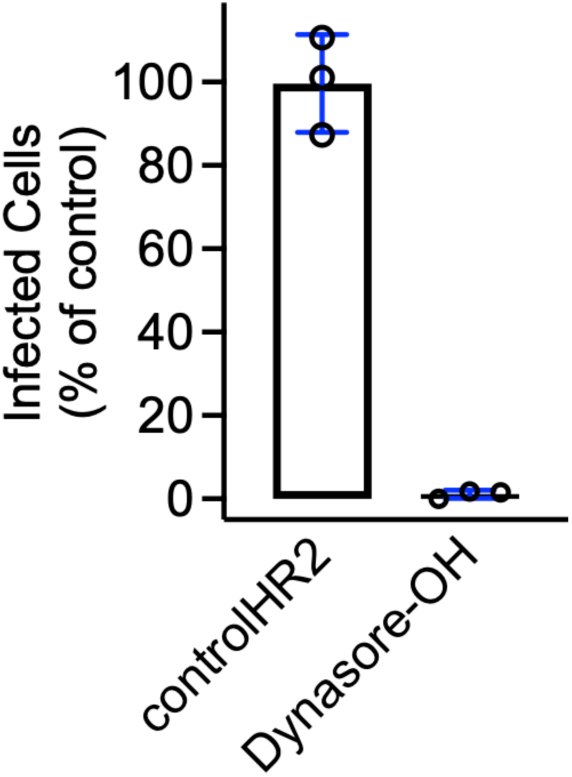
In the HR2 washout experiment, incubation of cells with controlHR2 had no effect on infection, while keeping dynasore-OH constant throughout the experiment completely inhibited infection.

**Fig. S5.**
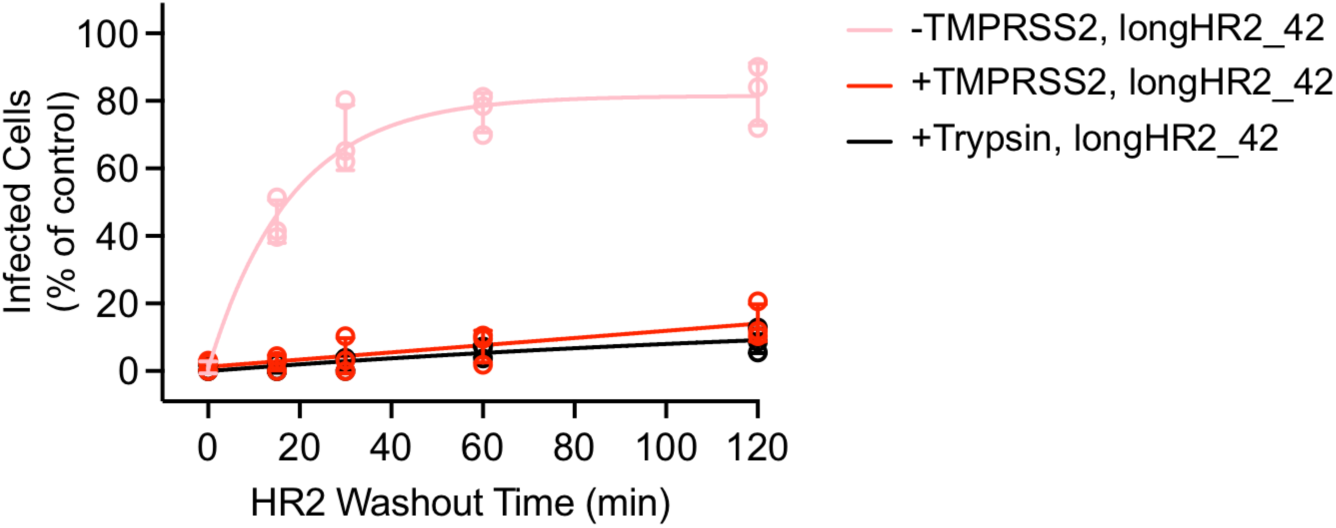
Trypsin treatment in TMPRSS2-negative VeroE6 cells shows similar inhibition as that of using TMPRSS2-expressing VeroE6 cells in the HR2 washout experiment. The VSV-SARS-CoV-2-Wuh chimera was treated with 1 µM trypsin for 30 min at 37°C followed by 10 µM Aprotinin, a trypsin inhibitor, before adding to the cells. Otherwise, the trypsin experiment was performed with the same condition as that of Fig. 3*E*.

**Fig. S6.**
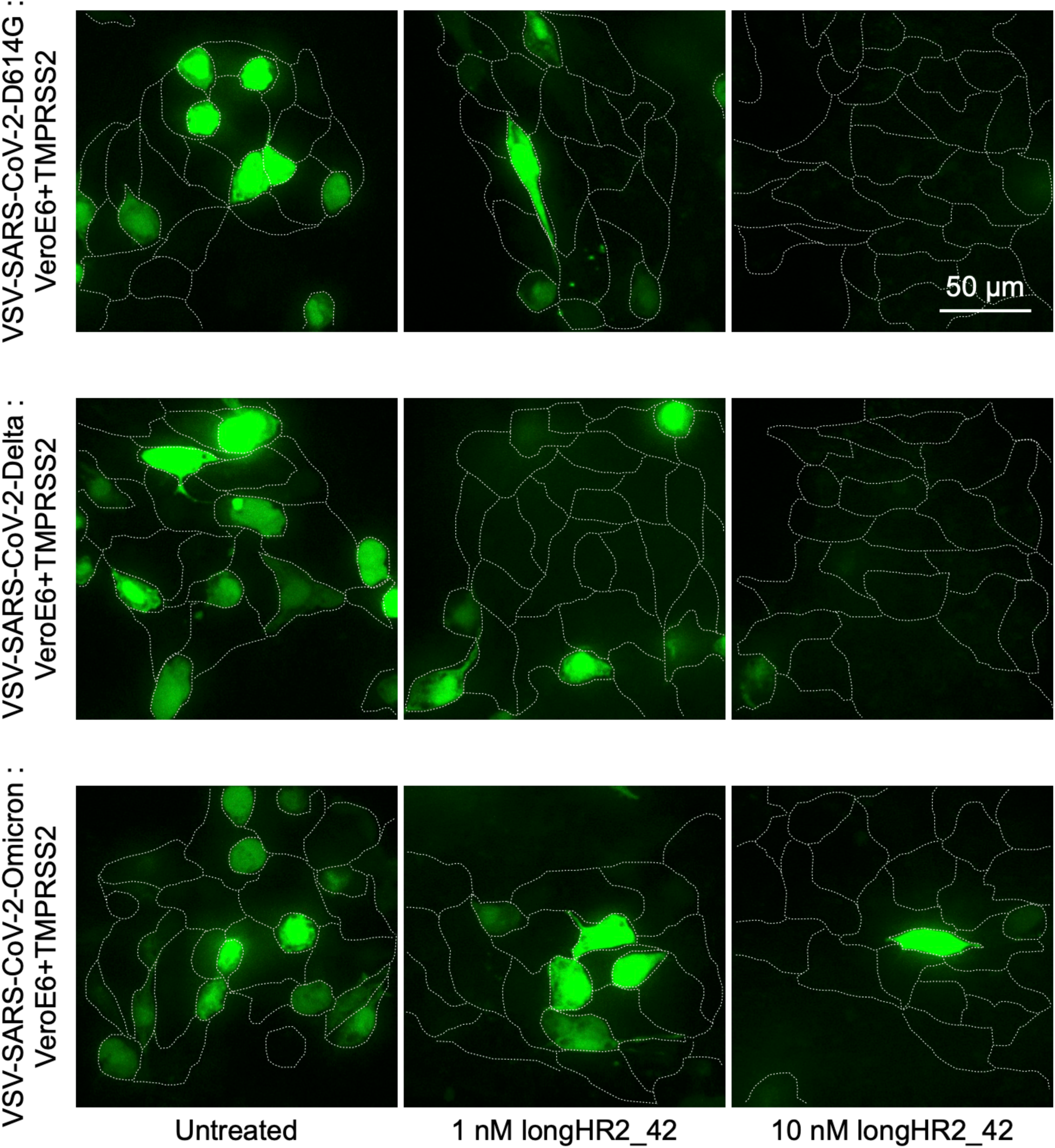
Example images of the inhibition of VSV-SARS-CoV-2-D614G, -Delta, and -Omicron infection by longHR2_42. Example images are maximum intensity projections of 20 µm with 1 µm z-plane spacing taken on a spinning disc-confocal with eGFP expression allowing for infection to be determined. Cell outlines were obtained from a WGA-Alexa647 membrane stain.

**Fig. S7.**
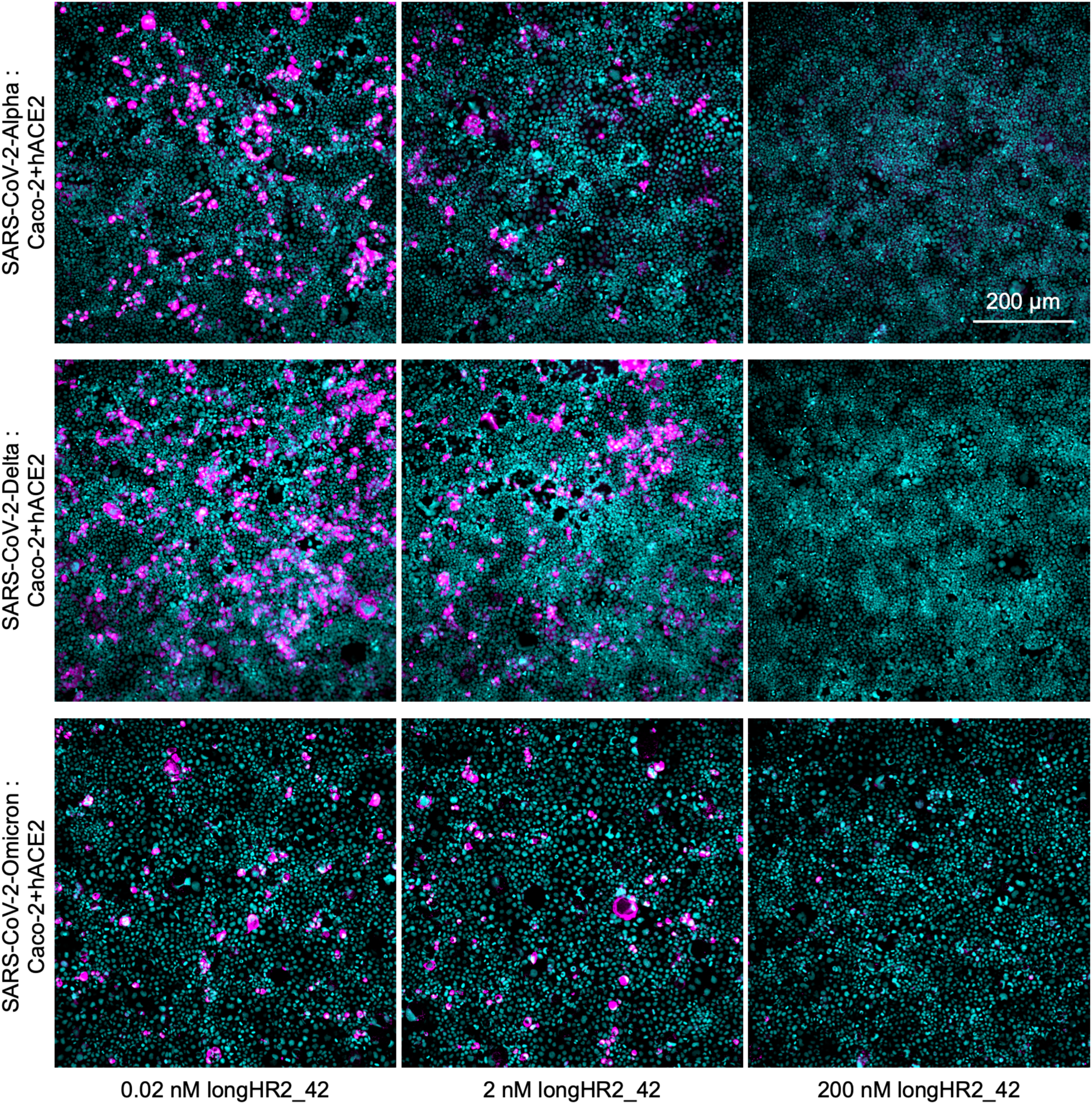
Example images of the inhibition of SARS-CoV-2-Alpha, -Delta, and -Omicron infection by longHR2_42. The infected cells are detected by an antibody specific for the viral N protein (magenta) and nuclei by Hoechst DNA dye (cyan).

**Fig. S8.**
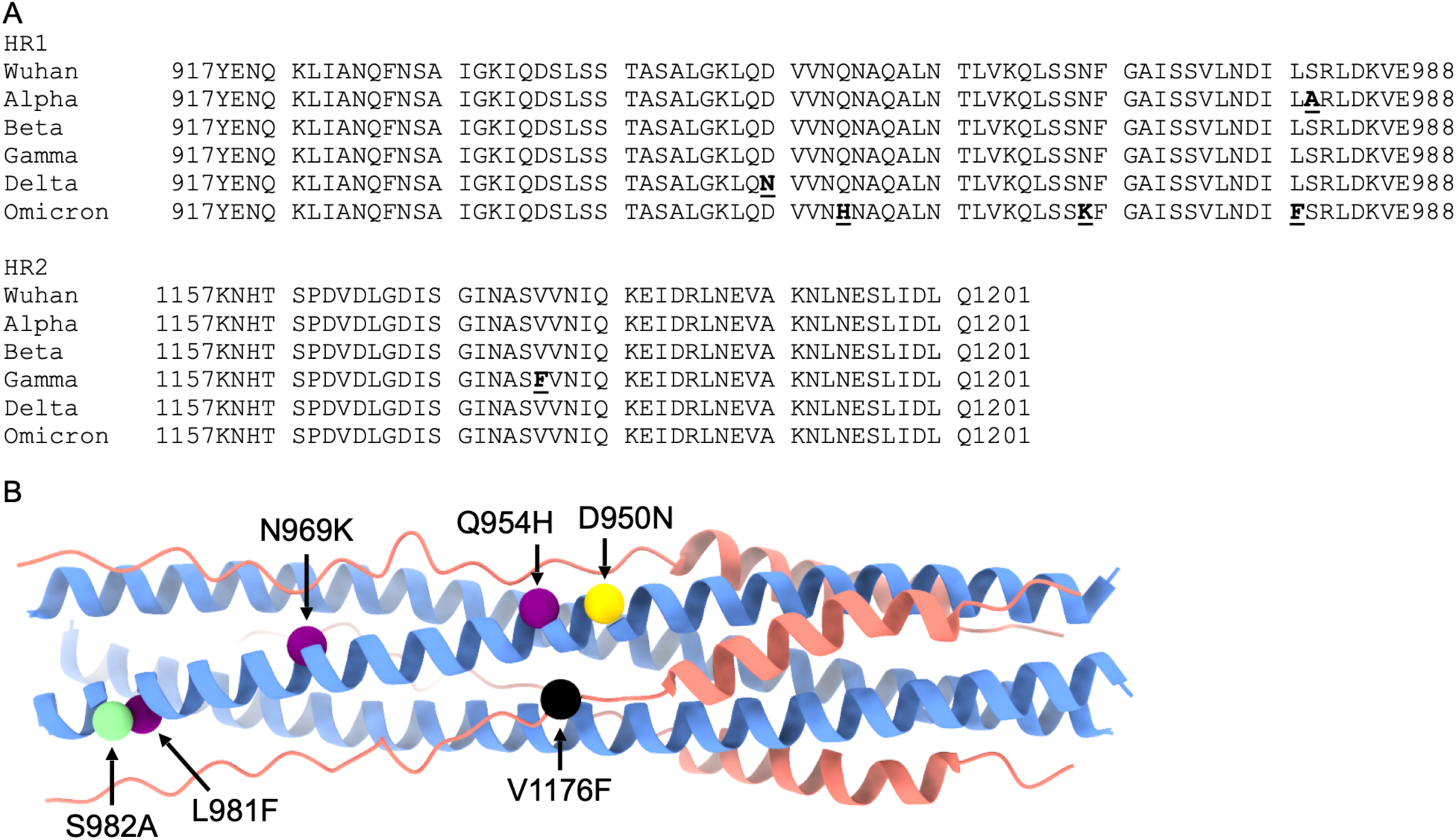
Mutations of SARS-CoV-2 variants of concern in the HR1HR2 region. (*A*) Sequence alignment of the HR1HR2 region of SARS-CoV-2 variants. (*B*) Locations of mutations in the 3D structure of HR1HR2 (PDB id 8czi). Color code: blue for HR1, red for HR2, green for Alpha, black for Gamma, yellow for Delta, and purple for Omicron.

**Fig. S9.**
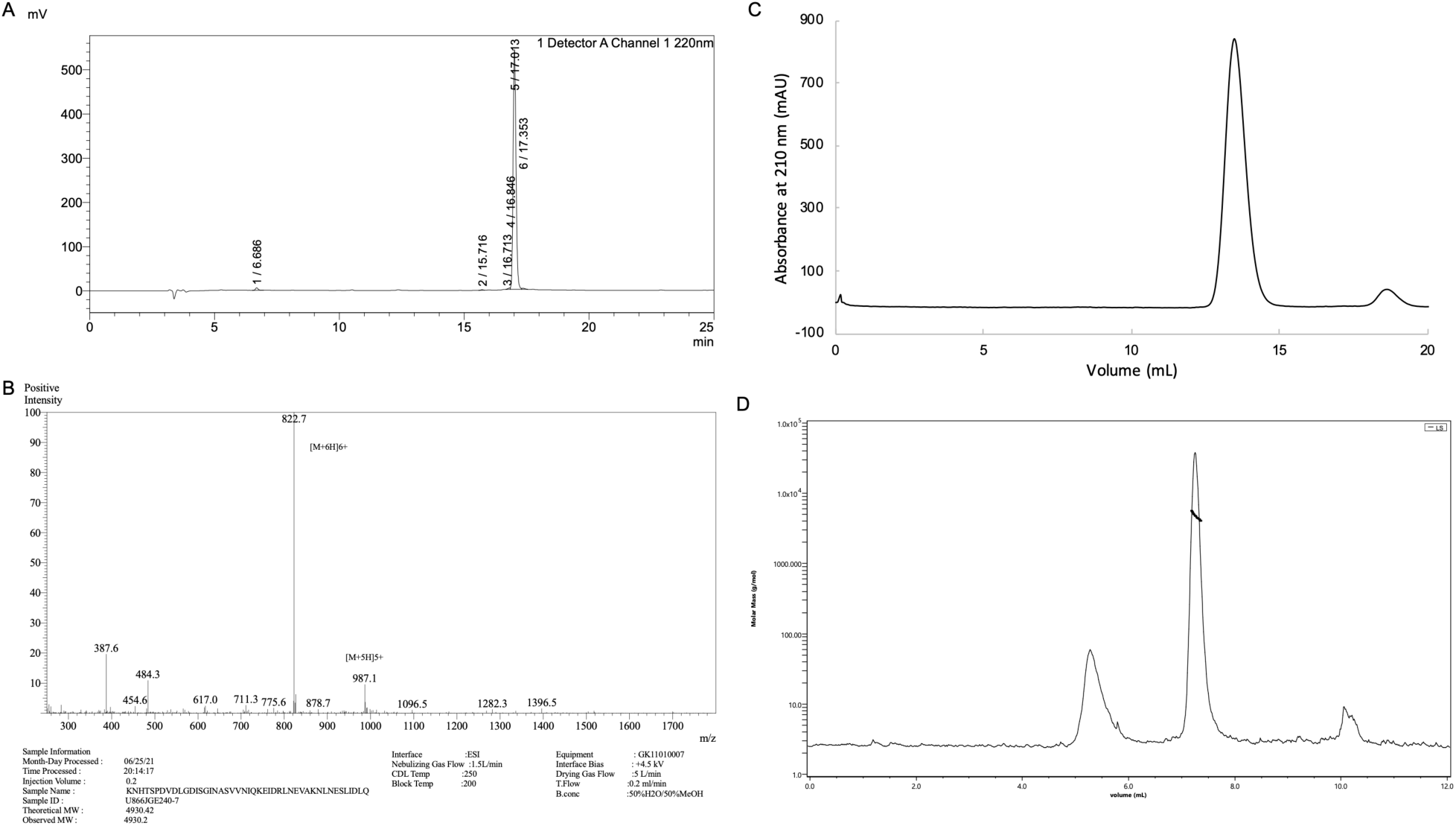
HPLC (*A*), LC-MS (*B*), SEC (*C*), and SEC-MALS (*D*) profiles of longHR2_45. The molecular weight measured by SEC-MALS is 4.8 kDa, close to 4.9 kDa, the theoretical monomer molecular weight.

**Fig. S10.**
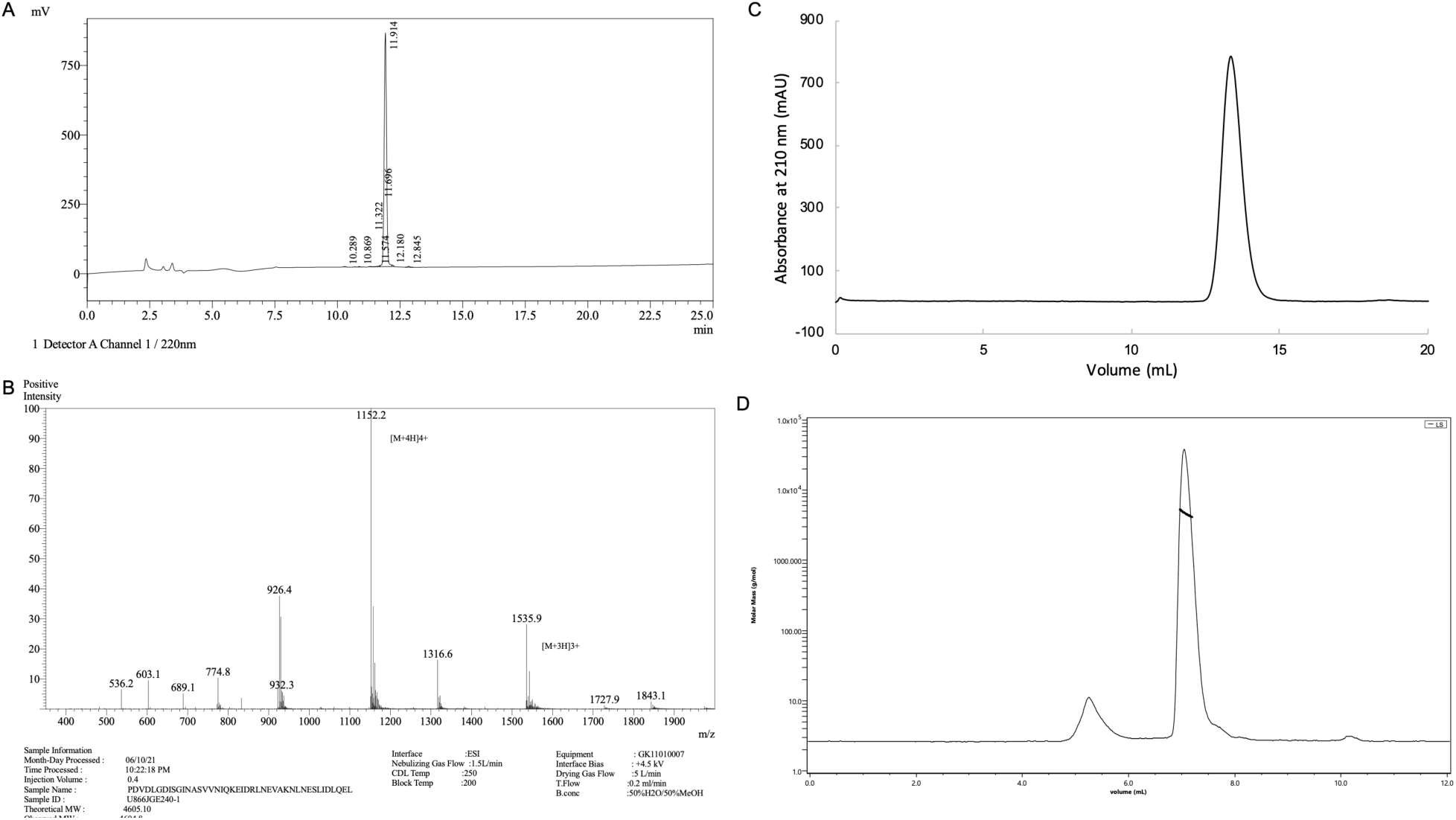
HPLC (*A*), LC-MS (*B*), SEC (*C*), and SEC-MALS (*D*) profiles of longHR2_42. The molecular weight measured by SEC-MALS is 4.7 kDa, close to 4.6 kDa, the theoretical monomer molecular weight.

**Fig. S11.**
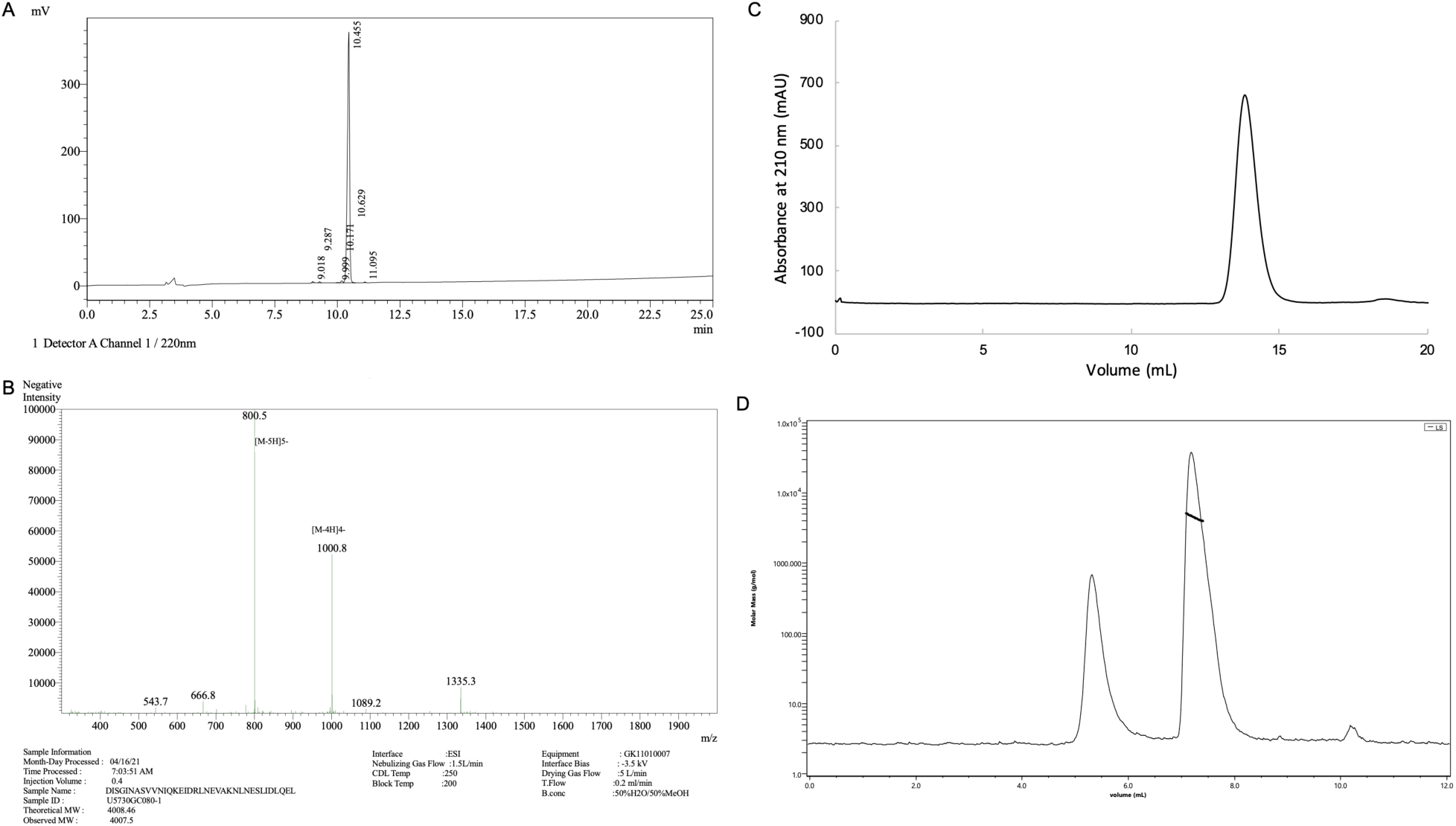
HPLC (*A*), LC-MS (*B*), SEC (*C*), and SEC-MALS (*D*) profiles of shortHR2. The molecular weight measured by SEC-MALS is 4.5 kDa, close to 4.0 kDa, the theoretical monomer molecular weight.

**Fig. S12.**
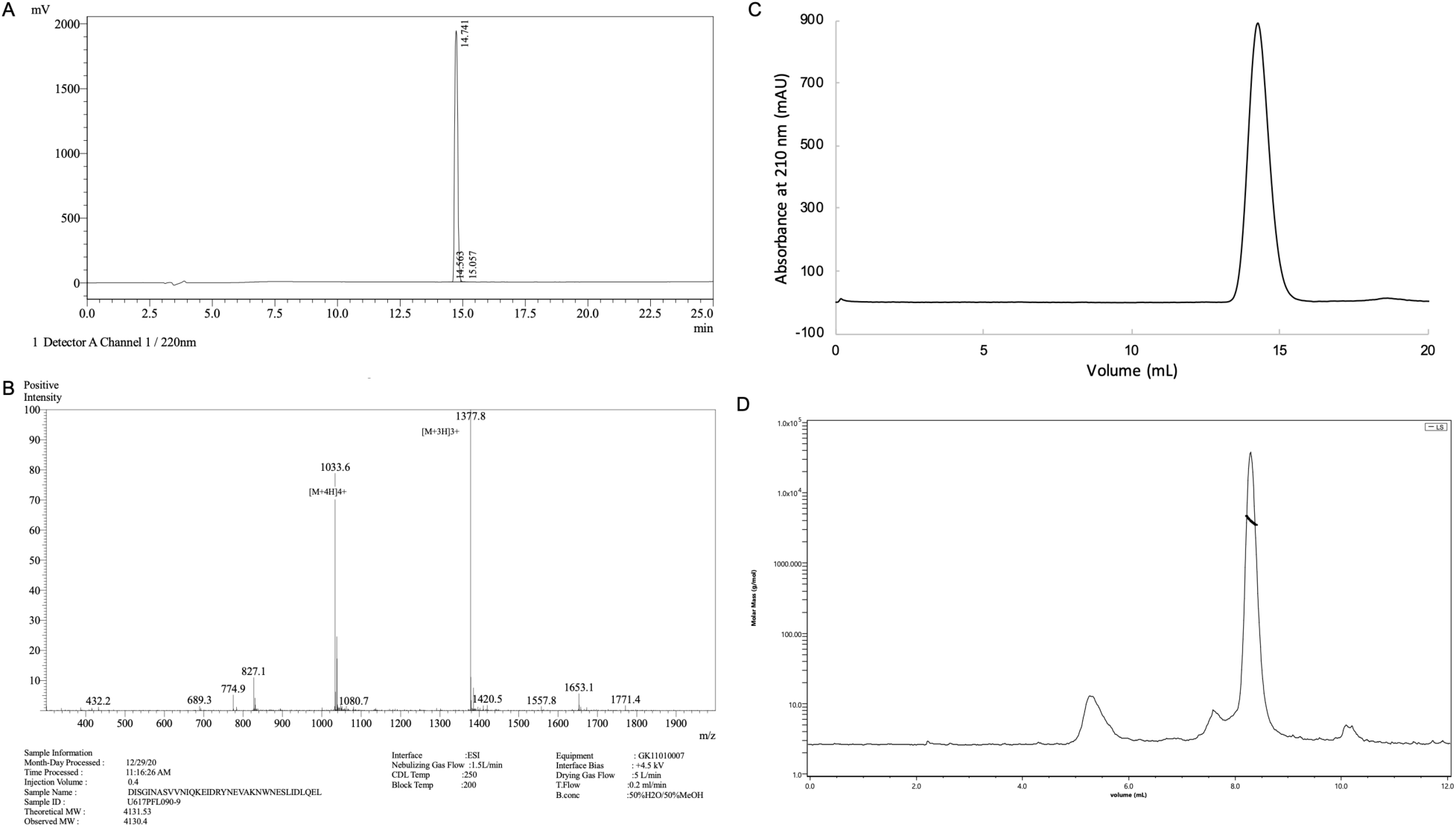
HPLC (*A*), LC-MS (*B*), SEC (*C*), and SEC-MALS (*D*) profiles of controlHR2. The molecular weight measured by SEC-MALS is 4.0 kDa, close to 4.1 kDa, the theoretical monomer molecular weight.

**Table S1.** Cryo-EM data collection, refinement, and model building.

|  |  |
| --- | --- |
|  | EMDB-27098, PDB 8czi |
| Microscope | Titan Krios |
| Voltage (kV) | 300 |
| Camera | Gatan K3 |
| Pixel size (Å) | 0.653 |
| Exposure time (s) | 1 |
| Number of frames per exposure | 40 |
| Total Dose (e<sup>-</sup>/Å<sup>2</sup>) | 47 |
| Number of movies | 18,846 |
| Defocus range (μm) | -2 to -0.3 |
| Number of particles | 751,443 |
| Resolution of final global refinement (0.143 FSC, Å) | 1.83 |
| Resolution of final local refinement (0.143 FSC, Å) | 2.22 |
| Bond RMSD (Å) | 0.004 |
| Angle RMSD (°) | 0.333 |
| Molprobity score | 1.37 |
| Clashscore, all atoms | 6.62 |
| Ramachandran favored (%) | 100 |
| Ramachandran allowed (%) | 0 |
| Ramachandran outliers (%) | 0 |
| Rotamer outliers (%) | 0 |
| Cβ outliers (%) | 0 |
| CaBLAM outliers (%) | 0 |

**Table S2.** Read counts of variants sequenced during mRNA display.

| Variant | 0 nM<br>input | 10 nM<br>input | 32 nM<br>input | 100 nM<br>input | 0 nM<br>eluate | 10 nM<br>eluate | 32 nM<br>eluate | 100 nM<br>eluate |
| --- | --- | --- | --- | --- | --- | --- | --- | --- |
| Scrambled<br>peptide | 65952 | 73405 | 52473 | 67097 | 2501 | 2765 | 449 | 657 |
| shortHR2 | 2105.19 | 2726.64 | 1687.21 | 2044.18 | 203.721 | 618.968 | 1812.75 | 2661.66 |
| longHR2_42 | 17948.6 | 25428.5 | 17014.4 | 22193.4 | 2156.4 | 10647.3 | 30756 | 44710.5 |

